# Hemispheric Dissociation in the Cognitive Control of Word Production

**DOI:** 10.64898/2026.06.12.731817

**Authors:** A. Yucel, A. K Martin

**Affiliations:** School of Psychology, University of Kent, Canterbury, UK; Kent Medway Medical School, Canterbury, UK

**Keywords:** focal tDCS, inferior frontal gyrus, word production, picture–word interference, picture description

## Abstract

This study investigated the hemispheric contributions of the inferior frontal gyrus (IFG) to word production across constrained and unconstrained tasks. In the present study, we used focal ring transcranial direct current stimulation (tDCS) in a sham-controlled, double-blind design, combined with a Picture–Word Interference (PWI) task and a picture description task. Fifty-four healthy young adults completed both tasks within the same stimulation sessions, receiving anodal stimulation over either the left or right IFG. Following task-specific data exclusions, 52 participants were included in each of the two task analyses. In the PWI task, categorically related distractors produced semantic interference and associatively related distractors facilitated naming in response times, consistent with previous findings. Stimulation did not affect response times. In error rates, the two stimulation sites modulated the associative effect in opposite directions. Anodal stimulation of the left IFG enhanced the associative advantage in error rates, whereas anodal stimulation of the right IFG produced the opposite pattern. In the picture description task, anodal stimulation of the right IFG increased speech rate, producing significant effects on words per minute, unpruned words per minute, and syllables per minute. No equivalent effects emerged in the left IFG group. Lexical diversity and utterance length were unaffected by stimulation. Taken together, these findings provide convergent evidence for a hemispheric dissociation in word production. The right IFG appears to contribute to the regulation of speech output and the control of distractor interference, whereas the left IFG appears to support the retrieval and selection of word meanings.

## 1. Introduction

Word production is a fundamental ability supported by a widespread network of brain regions, with the left perisylvian cortex playing a dominant role (Indefrey & Levelt, 2004; Hickok & Poeppel, 2007). Within this network, the left inferior frontal gyrus has been linked to word selection, semantic control and the regulation of competition between candidate words (Thompson-Schill et al., 1997; Schnur et al., 2009). However, a growing number of studies are showing that the right inferior frontal gyrus makes systematic contributions to language production in both healthy speakers and clinical populations. Accordingly, the assumption of the absolute dominance of the left hemisphere has been questioned (Turkeltaub et al., 2011; Hartwigsen & Saur, 2019). The nature of these contributions is not yet fully understood. Most importantly, it remains unclear whether the left IFG and the right IFG operate in parallel during word production or engage in functionally distinct cognitive processes. Causal inferences about relationships between brain and behaviour can be drawn using non-invasive brain stimulation techniques, such as transcranial direct current stimulation (Sellaro et al., 2016). We used focal ring tDCS to investigate whether the left IFG and right IFG support dissociable cognitive operations during word production. In the present study, two complementary picture-based paradigms were administered within the same experimental session.

Picture naming has been widely used to examine the cognitive mechanisms underlying word production. The picture–word interference (PWI) paradigm places naming under controlled conditions of lexical competition (Glaser & Düngelhoff, 1984). Participants name a target picture while attempting to ignore a superimposed distractor word whose semantic relationship to the target can be systematically varied. When the distractor belongs to the same semantic category as the target, naming latencies are slowed relative to an unrelated control, and this effect is termed ‘semantic interference’ (Schriefers et al., 1990; Damian & Martin, 1999). This effect has been attributed either to competition between target and distractor during lexical selection (Roelofs, 1992) or to a later, post-lexical process that excludes the distractor from production (Mahon et al., 2007). When the distractor is associatively related to the target, naming latencies are often sped up instead, an effect attributed to conceptual or lexical priming through spreading activation (Alario et al., 2000; Abdel Rahman & Melinger, 2009). These two effects emerge at distinct points in the time course of word production. Meta-analytic estimates indicate that conceptual processing occurs within the first 200 ms after picture onset, lemma selection between approximately 200 and 350 ms, and phonological encoding thereafter (Indefrey & Levelt, 2004; Indefrey, 2011). Their dissociable temporal profiles across stimulus-onset asynchronies have been taken as evidence that they engage at least partially separable stages of word production (La Heij et al., 1990; Sailor et al., 2009). Both effects are most often measured in naming latencies and only rarely in naming accuracy, which typically approaches the ceiling (Arrigoni et al., 2025). The neural basis of these effects has also been examined. Neuroimaging studies of the picture–word interference task have implicated the left posterior and middle temporal cortex in semantic interference (de Zubicaray et al., 2001). The contribution of the left IFG has been more debated. Lesion evidence points to a functional division. In one study, lesions to the left IFG increased categorical interference, while lesions to posterior temporal regions increased associative facilitation, with both effects evident in error rates (Pino et al., 2022). This dissociation points to a causal role for the left IFG in categorical competition. We designed our left IFG stimulation to target this process.

A picture description task provides an additional measure of word production under unconstrained conditions. In this paradigm, speakers describe a complex visual scene without time pressure, allowing assessment of speech rate, lexical diversity, and syntactic complexity in the absence of an explicit competing distractor (Wilson et al., 2010; Forbes-McKay & Venneri, 2005). The task is thought to engage processes related to message planning, lexical retrieval, and the regulation of speech output (Levelt et al., 1999). Neuroimaging indicates that connected speech production recruits a broader network than single-word naming. Describing a picture story engages the bilateral prefrontal cortex, including the inferior frontal gyrus of both hemispheres (Troiani et al., 2008). Narrative production also recruits regions beyond the classical language network, such as premotor and medial prefrontal areas (AbdulSabur et al., 2014). Therefore, the PWI paradigm and picture description place different demands on the word production system. The PWI paradigm measures the resolution of competition between target and distractor under controlled conditions, whereas picture description measures the regulation of output during continuous speech. Combining both paradigms allows us to test how each hemisphere contributes to word production under high and low levels of lexical competition.

One prominent account links the right IFG to response inhibition and cognitive control. Specifically, the right IFG has been described as a brake that can stop or slow an ongoing response (Aron et al., 2014). This brake can be engaged by external signals, such as a stop cue, or by internal goals. However, competing evidence suggests that the role of the right IFG is not inhibition alone. It may instead support the broader detection of salient or task-relevant information, with inhibition being one outcome of this wider function (Hampshire et al., 2010). Cognitive control of this kind is also relevant to language production. For example, speakers must select a target word while suppressing competing alternatives. This selection process has been linked to domain-general control mechanisms (Nozari & Novick, 2017). A separate line of research has linked the right hemisphere to semantic processing itself. Studies of language comprehension suggest that the two hemispheres process meaning in different ways. The left hemisphere supports focused activation of dominant meanings, whereas the right hemisphere activates a broader range of related concepts (Jung-Beeman, 2005; Beeman, 1993). This coarse form of semantic activation has been associated mainly with right temporal regions, although the right IFG may also contribute(Vigneau et al., 2011). However, whether these processes operate during word production, rather than comprehension, remains an open question. Together, these lines of work suggest that the right IFG could contribute to word production through both cognitive control and semantic context processing.

A second account links the right IFG to the regulation and monitoring of speech output. Models of speech motor control include a right-lateralised component. This component monitors and corrects ongoing speech using sensory feedback (Tourville & Guenther, 2011). The right IFG has also been shown to directly influence speech production. For example, the left IFG can be temporarily disrupted in healthy speakers. When this happens, activity in the right IFG increases, and this region provides a facilitatory drive to the left IFG (Hartwigsen et al., 2013). Importantly, responses become faster as the influence of the right IFG grows. This indicates an active rather than incidental contribution. This pattern fits a broader view of the right hemisphere as a latent part of the language network. Evidence from aphasia recovery suggests that right-hemisphere homologues of left language regions are recruited when the left hemisphere is damaged (Turkeltaub et al., 2011; Hartwigsen & Saur, 2019). On this view, these regions are not new additions but weakly active parts of a bilateral network that become more active under altered conditions. The right IFG may therefore contribute to word production through the monitoring of speech output, with a functional role that is normally masked by left-hemisphere dominance. Consistent with this regulatory role, modulating the right IFG has been shown to alter the speed of word production (Rosso et al., 2014). The left IFG, by contrast, has an established role in lexical selection and in syntactic processing (Thompson-Schill et al., 1997; Embick et al., 2000). During connected speech, these functions are likely to shape the diversity of words retrieved and the length of utterances, rather than the rate of output.

Testing these proposed roles requires a method that can isolate the contribution of a single region. Transcranial direct current stimulation modulates cortical excitability by passing a weak electrical current between electrodes placed on the scalp (Stagg & Nitsche, 2011). However, conventional tDCS uses two large electrodes, which spread the current across a wide cortical area. This diffuse current makes it difficult to attribute behavioural effects to a specific region. Focal ring tDCS addresses this limitation. It uses a small central electrode surrounded by a concentric ring, which confines the current to a more restricted area (Bortoletto et al., 2016). This precision is important for the present study, which compares stimulation of the left and right IFG. A direct comparison of the two regions is only meaningful if the current can be confined to each region in turn.

Previous brain stimulation studies of word production have focused almost exclusively on the left hemisphere and on single-word tasks. An early tDCS study of picture naming reported faster naming after anodal stimulation of left posterior temporal cortex (Sparing et al., 2008). Subsequent work targeted the left frontal cortex. For example, anodal tDCS over the left prefrontal cortex reduced semantic interference during picture naming (Wirth et al., 2011). Stimulation of the left IFG also reduced semantic interference, while stimulation of the left superior temporal gyrus increased it (Pisoni et al., 2012). However, these effects have not always been replicated. In a PWI study, anodal tDCS over the left IFG did not modulate semantic interference, although left middle temporal gyrus stimulation reduced associative facilitation (Henseler et al., 2014). Moreover, across four experiments using frontal and temporal tDCS, real stimulation did not differ from sham, even in conditions requiring the suppression of semantic interference (Westwood et al., 2017). The right IFG has been targeted only rarely. In one of the few such studies, cathodal stimulation of the right IFG sped picture naming in healthy speakers (Rosso et al., 2014). Anodal stimulation of the right IFG has been examined only in connected speech (Matar et al., 2020), not in picture naming or PWI. Taken together, this literature indicates that left frontal and temporal regions contribute to word production. However, the pattern of results is inconsistent, and the role of the right IFG remains largely unexplored. These studies also used conventional montages, which limit spatial precision. More recent work has applied focal ring tDCS to the IFG during word production (Yucel et al., 2025; Yucel & Martin, 2026).

This literature is also limited in the tasks it has used. Word production has been examined almost entirely through single-word tasks. Connected speech production has received far less attention. The few relevant tDCS studies have been conducted mainly in clinical populations. For example, anodal stimulation of the left IFG improved linguistic cohesion in chronic non-fluent aphasia (Marangolo et al., 2014). Anodal stimulation of the same region also reduced disfluency in adults who stutter (Chesters et al., 2018). A case study further reported possible benefits for connected speech production in progressive supranuclear palsy (Madden et al., 2019). In healthy speakers, connected speech has rarely been examined. Anodal tDCS over the left and right IFG improved discourse production in healthy older adults, although the two sites did not differ significantly (Matar et al., 2020). That study used a small sample, a within-subjects design, and a conventional electrode montage. No tDCS study has examined speech rate during picture description in healthy adults. More importantly, no study has examined the PWI paradigm and connected speech within the same design. It therefore remains unknown whether stimulation of the left and right IFG produces convergent effects across constrained and unconstrained word production.

Taken together, the left and right IFG may support dissociable operations during word production. However, no study has tested this across both constrained and unconstrained tasks. The current study investigated this question in healthy adults, using two picture-based paradigms within a single session. For the PWI task, anodal stimulation of the left IFG was expected to reduce semantic interference without affecting associative facilitation. Anodal stimulation of the right IFG was expected to show the reverse pattern, modifying associative facilitation without affecting semantic interference. Anodal stimulation was also expected to reduce naming errors in both conditions. Increasing the excitability of the targeted region should support more efficient retrieval and selection, regardless of distractor type. For connected speech, anodal right IFG stimulation was expected to increase speech rate, whereas anodal left IFG stimulation was expected to increase lexical diversity and utterance length. At a broader level, these predictions reflect a proposed division of labour between the two regions. The left IFG was expected to contribute to the retrieval and selection of word meanings, and the right IFG to the regulation of speech output.

## 2. Method

### 2.1. Participants

Fifty-four participants took part in the study and were assigned to one of two stimulation groups: 26 to the left IFG group and 28 to the right IFG group. The left IFG group consisted of 20 females and 6 males, and the right IFG group consisted of 20 females and 8 males. The groups were comparable in gender distribution, χ*²*(1, N = 54) = 0.21, *p* = .645. Mean age did not differ between the left IFG group (*M* = 19.35 years, *SD* = 0.85) and the right IFG group (*M* = 20.14 years, *SD* = 4.33), t(52) = −0.92, *p* = .361, *d* = −0.25. Each participant completed both the picture–word interference (PWI) task and the picture description task within the same two stimulation sessions. Owing to task-specific data exclusions, two right IFG participants were excluded from the picture–word interference study due to technical recording problems, and two right IFG participants were excluded from the picture description study for not meeting the minimum word count required for analysis. Therefore, each study included 52 participants (26 with left IFG, 26 with right IFG), with 50 participants contributing data to both.

Participants were recruited from the University of Kent student population and received course credit upon completion. All were right-handed, spoke English as a first language, were neurologically healthy, and had no prior experience with transcranial direct current stimulation (tDCS). They reported no history of neurological or psychiatric conditions and had normal or corrected-to-normal vision. Prior to participation, all individuals were screened against standard tDCS safety criteria. Exclusion criteria included a history of neurological disease, a current or past psychiatric diagnosis requiring treatment, metal implants or medical devices incompatible with tDCS, the use of medications known to influence cortical excitability (e.g., antidepressants, anxiolytics), and vascular or metabolic conditions known to affect cognition. Two individuals were excluded at screening, one for a history of epilepsy and one for antidepressant use for major depression. All participants gave written informed consent prior to participation. Ethical approval was obtained from the Human Research Ethics Committee at the School of Psychology, University of Kent (ID: 202216587800677857).

### 2.2. Design and Procedure

A double-blind, sham-controlled, mixed design was used, with stimulation condition (sham, anodal) as a within-subjects factor and stimulation region (left IFG, right IFG) as a between-subjects factor. Each participant attended two experimental sessions, separated by at least 72 hours. One session involved active anodal tDCS, and the other involved sham stimulation. The order of the two stimulation conditions was counterbalanced across participants. Assignment to the left or right IFG group was fixed at recruitment and remained constant across both sessions. All testing took place in the brain stimulation laboratories at the School of Psychology, University of Kent.

The two sessions followed an identical structure. Mood was assessed at the start of each session, before stimulation. Stimulation then began, and both experimental tasks were conducted online during the 20-minute stimulation period. Participants completed the picture–word interference task first, followed by the picture description task. Mood and adverse effects were assessed at the end of each session. The task-specific procedures are described separately for each study below.

### 2.3. Focal Transcranial Direct Current Stimulation

Transcranial direct current stimulation (tDCS) was delivered using a battery-operated DC stimulator (DC-Stimulator Plus, NeuroConn, Ilmenau, Germany) in a double-blind, sham-controlled design. A focal electrode montage was employed, consisting of a small circular centre electrode (2.5 cm diameter) surrounded by a ring-shaped return electrode (inner diameter: 7.5 cm; outer diameter: 10 cm). This configuration was selected to enhance spatial focus and increase current density over the target cortical region.

The centre electrode was placed over the inferior frontal gyrus following the international 10–20 EEG system, corresponding to FC5 for the left IFG and FC6 for the right IFG. A ring-shaped return electrode was positioned around the centre electrode to create a more focal stimulation setup. To examine the distribution of the induced electric field, current-flow simulations were performed using SimNIBS (Thielscher et al., 2015). These simulations were conducted on a standard MNI152 T1-weighted brain template with 1 mm resolution. The resulting electric field was visualised, and the normal component of the electric field is shown in Figure 1. This component is considered most relevant for modulating neuronal excitability (Radman et al., 2007).

**Fig. 1.**
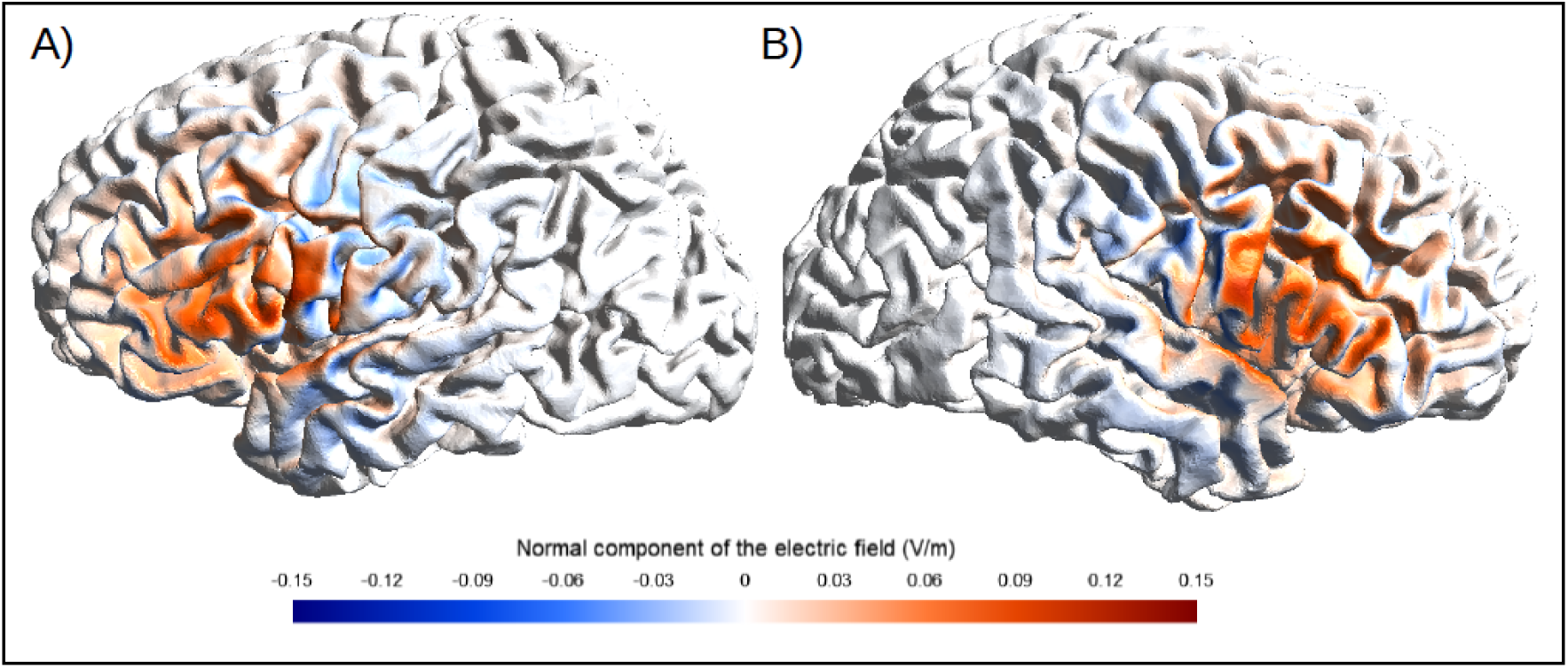
Simulated electric field under anodal stimulation for the two target sites. (A) Left inferior frontal gyrus. (B) Right inferior frontal gyrus. Note. Surface renderings show the distribution and intensity of the normal component of the simulated electric field (V/m).

In both stimulation conditions, current intensity was ramped up to 1 mA over 5 seconds. In the anodal condition, stimulation was maintained at 1 mA for 20 minutes, then ramped down over 5 seconds. In the sham condition, current was applied for approximately 40 seconds before ramping down, producing initial cutaneous sensations comparable to active stimulation and thereby supporting effective participant blinding. Stimulation was administered using the device’s study mode, which employs pre-programmed stimulation codes to determine stimulation conditions. These codes were assigned by a researcher not involved in data collection, ensuring that both participants and experimenters remained blind to the stimulation condition.

### 2.4. Mood, Adverse Effects, and Blinding

Mood was assessed before and after each session using the Visual Analogue Mood Scale developed by Folstein and Luria (1973). On this scale, participants rate their current emotional state on a computerised slider ranging from 0 to 100. Higher values indicate a stronger feeling. Ratings were obtained for eight states: fear, confusion, sadness, anger, energy, fatigue, happiness, and tension. For each state, a mood change score was calculated by subtracting the pre-stimulation rating from the post-stimulation rating. The eight states were then grouped into a positive set and a negative set. The positive set consisted of energy and happiness. The negative set consisted of fear, confusion, sadness, anger, fatigue, and tension. A positive and a negative mood score were then calculated by adding the change scores within each set. These two scores were analysed separately.

Adverse effects of stimulation were assessed at the end of each session. A standardised, computerised self-report questionnaire adapted from Brunoni and colleagues (2011) was used. Participants rated each sensation on a four-point scale ranging from absent to severe. The questionnaire covered sensations commonly linked to tDCS, including headache, neck discomfort, scalp sensations, tingling, itching, burning, skin redness, sleepiness, and difficulty concentrating. A total adverse effects score was then calculated for each session by adding the ratings across all items.

Blinding was assessed at the end of the second session. Participants were asked to indicate which of their two sessions they believed had involved active stimulation.

### 2.5. Statistical Analyses

All analyses were performed in JASP (version 0.18.3). An alpha level of .05 was applied throughout, and partial eta squared (η*²*□) was used as the measure of effect size. Response times and error rates were analysed using mixed-design analyses of variance (ANOVAs). Separate 2 × 2 × 2 ANOVAs were conducted for the category and association conditions, with Stimulation (sham, anodal) and Interference (related, unrelated) as within-subjects factors and Region (left IFG, right IFG) as a between-subjects factor. Significant interactions involving Region were followed up with separate 2 × 2 ANOVAs within each region.

For the picture description task, each of the five speech rate measures was analysed in a separate 2 × 2 mixed-design ANOVA, with Stimulation as a within-subjects factor and Region as a between-subjects factor. Significant Stimulation × Region interactions were followed up with one-way repeated-measures ANOVAs within each region.

Adverse effects, positive mood scores, and negative mood scores were each analysed in a separate 2 × 2 mixed-design ANOVA, with Stimulation as a within-subjects factor and Region as a between-subjects factor. Significant Stimulation × Region interactions were followed up within each region. Blinding was assessed using a binomial test against the chance level of 50%, together with a chi-square test of association between Region and blinding accuracy.

## 3. Results

### 3.1. Adverse Effects, Mood, and Blinding

All participants completed both the PWI and picture description tasks within the same stimulation sessions. Adverse effects, mood, and blinding were measured at the session level and applied to both Study 1 and Study 2. These measures are reported once for the combined cohort, rather than separately within each study. A 2 × 2 mixed-design ANOVA revealed a main effect of Stimulation, *F*(1, 51) = 6.91, *p* = .011, η*²*□ = .119. The main effect of Region did not reach significance, *F*(1, 51) = 3.98, *p* = .051, η*²*□ = .072, but a significant Stimulation × Region interaction was observed, *F*(1, 51) = 5.81, *p* = .020, η*²*□ = .102. Follow-up analyses indicated that the interaction was driven by the right IFG group, in which anodal stimulation elicited higher adverse effects than sham, *F*(1, 26) = 8.64, *p* = .007, η*²*□ = .250. In the left IFG group, adverse effects did not differ between stimulation conditions, *F*(1, 25) = 0.05, *p* = .828, η*²*□ = .002.

There was no significant effect of Stimulation on either positive mood, *F*(1, 49) = 0.10, *p* = .757, η*²*□ = .002, or negative mood, *F*(1, 49) = 2.35, *p* = .132, η*²*□ = .046. No main effect of Region or Stimulation × Region interaction was observed for either measure (all *p*s > .20), indicating that mood ratings were comparable across stimulation conditions and regions.

**Table 1.** Descriptive statistics of adverse effects and mood change (positive and negative item sets) in the stimulation sessions.

|  | Left IFG |  | Right IFG |  |
| --- | --- | --- | --- | --- |
|  | Sham | Anodal | Sham | Anodal |
|  | Mean | Mean | Mean | Mean |
|  | (SD) | (SD) | (SD) | (SD) |
| Adverse effects | 2.85 | 2.92 | 3.59 | 5.37 |
|  | (2.36) | (2.35) | (3.56) | (4.06) |
| VAMS positive | -6.65 | -3.04 | 5.04 | -1.08 |
|  | (22.75) | (23.71) | (35.59) | (19.62) |
| VAMS negative | -14.08 | -24.04 | -21.08 | -37.56 |
|  | (38.60) | (45.09) | (50.96) | (63.38) |

*Note.* SD = standard deviation; IFG = inferior frontal gyrus; VAMS = Visual Analogue Mood Scale

A binomial test indicated that blinding was effective. The proportion of correct guesses (33 of 54, 61.1%) did not differ significantly from the chance level of 50% (p = .134). A chi-square test revealed no association between Region and blinding accuracy, χ*²*(1, N = 54) = 0.39, *p* = .535, indicating that blinding did not differ between groups. The complete output for adverse effects, mood changes, and blinding is reported in Supplementary Table S3.

### 3.2. Study 1

#### 3.2.1. Picture–Word Interference Task

During each experimental session, participants completed a Picture–Word Interference (PWI) picture naming task while receiving either anodal or sham tDCS. Four distractor conditions were included: categorically related, associatively related, unrelated controls for categorical distractors, and unrelated controls for associative distractors. The unrelated control conditions were constructed by re-pairing distractor words with target pictures that bore no semantic or associative relationship to them.

At the start of each session, participants completed a short practice block during which they were instructed to name each picture as quickly and accurately as possible while ignoring the distractor word. The practice block was followed by the main PWI naming task. The main task consisted of 160 trials per session, organised into four successive blocks and separated by short breaks. Two blocks contained categorical trials, and two blocks contained associative trials. Each block comprised 40 trials, including 20 related and 20 unrelated picture–distractor pairs. Therefore, the full task included 80 categorical trials and 80 associative trials. Categorical blocks were always presented before associative blocks, and block order was not counterbalanced. Within each block, related and unrelated trials were presented in pseudorandomised order.

Stimulus presentation and data collection were controlled using PsychoPy (Peirce et al., 2019). Naming responses were recorded using a voice key triggered at speech onset, and response times were measured with millisecond precision using the Black Box Toolkit. All trials were verified offline through manual inspection of audio recordings. Trials in which the voice key was triggered by non-speech sounds, such as coughs or microphone noise, were identified during this inspection, and the response time was corrected based on the true speech onset in the recording. An auditory cue marked the beginning of each trial. Errors were defined as incorrect naming responses, hesitations, or disfluencies, and only correct responses were included in the response time analyses.

#### 3.2.2. Trial Structure

Each trial began with a 2000 ms blank screen, followed by a 200 ms auditory cue. A distractor word was then presented at a stimulus-onset asynchrony (SOA) relative to picture onset, with negative SOAs indicating that the distractor preceded the target picture. Categorical distractors were presented at an SOA of −300 ms and associative distractors at an SOA of −100 ms. The target picture was displayed for 3000 ms, and the distractor word remained on screen throughout picture presentation. The trial structure is illustrated in Figure 2.

**Fig. 2.**
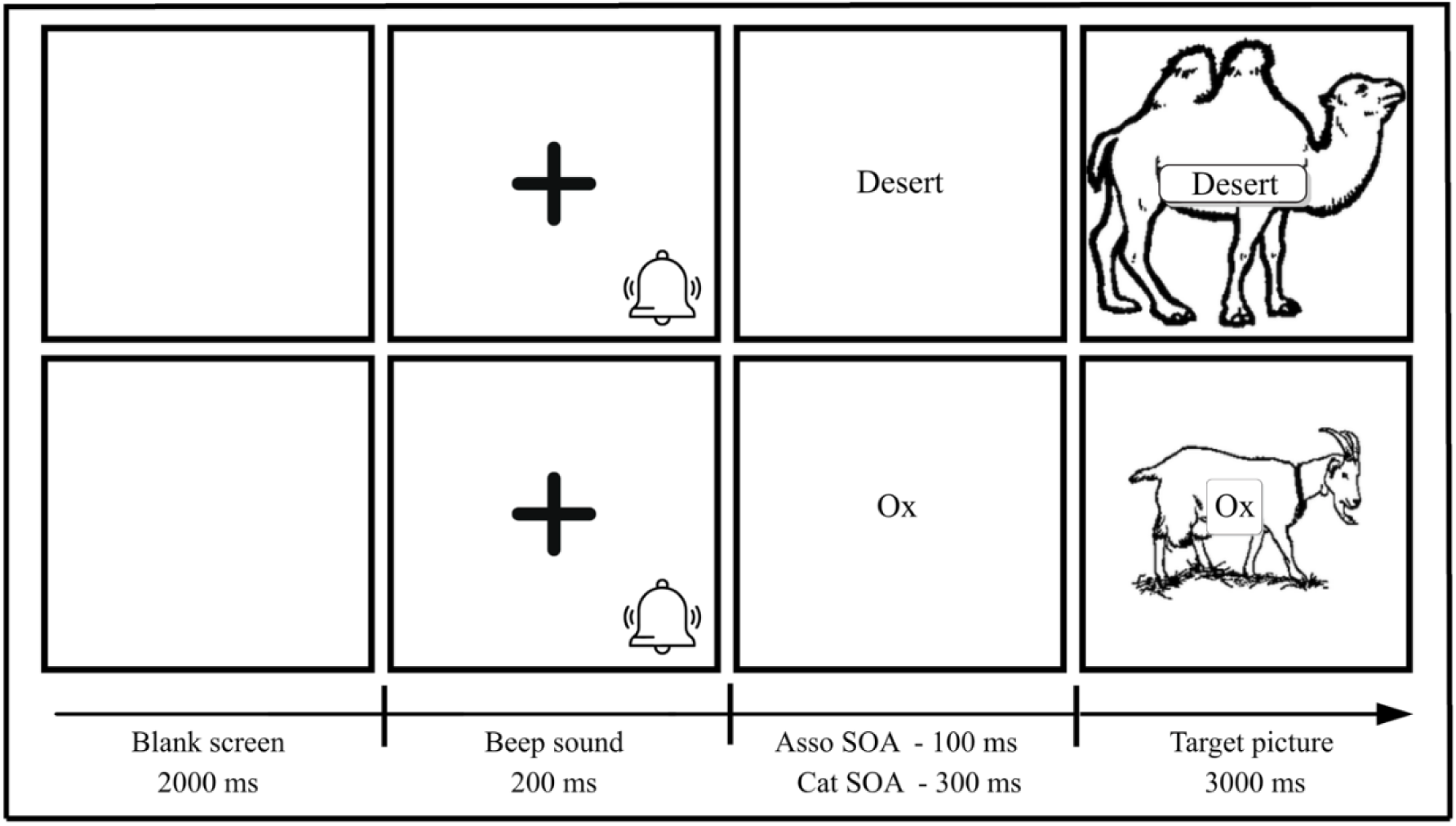
Trial structure of the picture–word interference (PWI) paradigm. *Note.* The upper row illustrates an associative trial and the lower row a categorical trial.

The categorical SOA of −300 ms was selected to allow pre-activation of the distractor’s semantic neighbours prior to the target’s lemma selection window. Categorical interference is thought to arise from competition between target and distractor lemmas in the same semantic category. This competition is mediated by lateral inhibition between co-activated representations (Levelt et al., 1999). For lateral inhibition to influence lemma selection, activation must first spread across the semantic network. An SOA of −300 ms provides sufficient time for this process, ensuring that related category members are active prior to the target’s lemma selection window at 200–350 ms after picture onset (Indefrey & Levelt, 2004). This timing is consistent with the original report by Glaser and Düngelhoff (1984), who observed a 69-ms categorical interference effect at this SOA, and with subsequent replications in healthy speakers (Hashimoto & Thompson, 2010; Sailor et al., 2009).

The associative SOA of −100 ms was selected to align the distractor’s peak activation with the target’s lemma selection window. The target’s lemma selection window has been estimated at approximately 200–350 ms after picture onset (Indefrey & Levelt, 2004; Indefrey, 2011). For written distractors, peak activation occurs at approximately 270–310 ms after the distractor itself appears on screen (Hauk et al., 2006; Krott et al., 2019). A distractor presented 100 ms before the picture would therefore reach its peak at approximately 170–210 ms after picture onset. This coincides with the onset of the target’s lemma selection window, which is the stage at which stimulation is most likely to modulate lexical access (Levelt et al., 1999; Roelofs, 1992).

A single SOA was used for each condition rather than a common SOA across both. The temporal profiles of categorical and associative effects in the picture–word interference literature do not converge on a single optimal value (Alario et al., 2000; Sailor et al., 2009). The two SOAs were held constant within their respective conditions, so that the categorical and associative effects could each be assessed at a single, fixed timing parameter.

#### 3.2.3. Stimuli

Pictures were selected from the International Picture Naming Project database (Szekely et al., 2004). Stimuli across conditions were matched on name agreement, number of syllables, number of letters, word frequency, and visual complexity, to ensure comparable difficulty across the associative, categorical, and unrelated control conditions (see Table 2). Categorically related distractors were selected using WordNet (Princeton University, 2010), whereas associatively related distractors were selected from the University of South Florida Free Association Norms (Nelson et al., 2004). Distractor word sets were also matched across conditions on number of syllables, number of letters, and lemma frequency. To maintain a clear separation between distractor conditions, categorically related distractors were selected without associative links to the target pictures, whereas associative distractors were chosen to avoid belonging to the same semantic category as the targets. Unrelated distractor–target pairs were constructed to ensure the absence of both semantic and associative relationships. Each target picture was presented only once per participant and assigned to a single distractor condition, whereas distractor words were reused across conditions by being paired with different target pictures.

**Table 2.** Characteristics of Stimuli.

| Feature | Associative | Unrelated<br>(Associative) | Categorical | Unrelated<br>(Categorical) | *p* |
| --- | --- | --- | --- | --- | --- |
| Name agreement | 0.86<br>(± 0.18) | 0.86<br>(± 0.15) | 0.85<br>(± 0.15) | 0.86<br>(± 0.15) | .988 |
| Number of syllables | 1.40<br>(± 0.50) | 1.37<br>(± 0.55) | 1.38<br>(± 0.59) | 1.50<br>(± 0.60) | .713 |
| Number of letters | 5.20<br>(± 1.47) | 4.94<br>(± 1.55) | 4.67<br>(± 1.16) | 5.03<br>(± 1.40) | .406 |
| Word frequency | 3.27<br>(± 1.04) | 3.37<br>(± 1.13) | 3.24<br>(± 1.26) | 3.19<br>(± 1.36) | .931 |
| Visual complexity | 17777<br>(± 10400) | 15889<br>(± 8438) | 15256<br>(± 6976) | 16547<br>(± 9171) | .624 |

Note. All values represent means (± standard deviations). *p*-values reflect omnibus comparisons across the four conditions. Picture stimuli characteristics were obtained from the International Picture Naming Project (https://crlexperiments.ucsd.edu/ipnp/).

#### 3.2.4. Results

##### 3.2.4.1. Response Times

###### Category Condition

Naming latencies in the category condition were analysed using a 2 × 2 × 2 mixed-design ANOVA with the factors Stimulation (anodal vs sham), Interference (categorically related vs unrelated), and Region (left IFG vs right IFG). Descriptive statistics are presented in Table 3. The complete ANOVA output for all picture–word interference analyses (response times and error rates) is reported in Supplementary Table S1. Naming latencies were significantly slower for categorically related compared to unrelated distractors, reflecting a semantic interference effect, *F*(1, 50) = 4.96, *p* = .030, η*²*□ = .090. There was no main effect of Stimulation, *F*(1, 50) = 2.74, *p* = .104, η*²*□ = .052, and no Stimulation × Region interaction, *F*(1, 50) = 0.32, *p* = .574, η*²*□ = .006. There was no interaction between Stimulation and Interference, *F*(1, 50) = 0.24, *p* = .629, η*²*□ = .005, and no three-way interaction between Stimulation, Interference, and Region, *F*(1, 50) = 0.30, *p* = .584, η*²*□ = .006, indicating that the semantic interference effect was present overall but was not modulated by stimulation or region. There was no main effect of Region, *F*(1, 50) = 0.94, *p* = .336, η*²*□ = .019.

**Table 3.** Descriptive statistics of naming performance (response times in milliseconds) in the PWI task.

|  | Left IFG |  | Right IFG |  |
| --- | --- | --- | --- | --- |
|  | Sham | Anodal | Sham | Anodal |
|  | Mean | Mean | Mean | Mean |
|  | (SD) | (SD) | (SD) | (SD) |
| Categorical | 1078 | 1050 | 1032 | 990 |
|  | (184) | (146) | (207) | (177) |
| Categorical Unrelated | 1037 | 1023 | 1021 | 978 |
|  | (163) | (150) | (202) | (159) |
| Associative | 1133 | 1118 | 1113 | 1076 |
|  | (176) | (148) | (204) | (214) |
| Associative Unrelated | 1181 | 1156 | 1126 | 1111 |
|  | (191) | (183) | (210) | (232) |

Note. Values are means, with standard deviations in parentheses. IFG = inferior frontal gyrus; SD = standard deviation.

###### Association Condition

Naming latencies in the association condition were analysed using the same model. Naming latencies were significantly faster for associatively related compared to unrelated distractors, reflecting an associative facilitation effect, *F*(1, 50) = 12.10, *p* = .001, η*²*□ = .195. There was no main effect of Stimulation, *F*(1, 50) = 1.38, *p* = .246, η*²*□ = .027, and no Stimulation × Region interaction, *F*(1, 50) = 0.03, *p* = .870, η*²*□ < .001. There was no interaction between Stimulation and Interference, *F*(1, 50) = 0.07, *p* = .787, η*²*□ = .001, and no three-way interaction between Stimulation, Interference, and Region, *F*(1, 50) = 0.59, *p* = .447, η*²*□ = .012, indicating that the associative facilitation effect was present overall but was not modulated by stimulation or region. There was no main effect of Region, *F*(1, 50) = 0.68, *p* = .412, η*²*□ = .013.

##### 3.2.4.2. Error Rates

###### Category Condition

Error rates in the category condition were analysed using a 2 × 2 × 2 mixed-design ANOVA with the factors Stimulation (anodal vs sham), Interference (categorically related vs unrelated), and Region (left IFG vs right IFG). There was a significant main effect of Interference, *F*(1, 50) = 12.86, *p* < .001, η*²*□ = .205, with fewer errors in related compared to unrelated trials. There was no main effect of Stimulation, *F*(1, 50) = 0.02, *p* = .877, η*²*□ < .001, and no Stimulation × Region interaction, *F*(1, 50) = 0.15, *p* = .698, η*²*□ = .003. There was no interaction between Stimulation and Interference, *F*(1, 50) = 0.53, *p* = .471, η*²*□ = .010, and no three-way interaction between Stimulation, Interference, and Region, *F*(1, 50) = 0.13, *p* = .718, η*²*□ = .003. There was no main effect of Region, *F*(1, 50) = 0.64, *p* = .429, η*²*□ = .013.

**Table 4.** Descriptive statistics of naming accuracy (error rates, %) in the PWI task.

|  | Left IFG |  | Right IFG |  |
| --- | --- | --- | --- | --- |
|  | Sham | Anodal | Sham | Anodal |
|  | Mean | Mean | Mean | Mean |
|  | (SD) | (SD) | (SD) | (SD) |
| Category | 5.769 | 5.769 | 5.385 | 6.154 |
|  | (5.512) | (5.948) | (4.165) | (5.578) |
| Category Unrelated | 7.308 | 7.019 | 8.846 | 8.750 |
|  | (3.737) | (5.149) | (5.841) | (4.016) |
| Associative | 4.712 | 4.038 | 5.000 | 5.769 |
|  | (3.628) | (3.813) | (5.339) | (5.906) |
| Associative Unrelated | 5.481 | 6.250 | 5.962 | 4.904 |
|  | (5.197) | (4.257) | (4.532) | (4.152) |

Note. Values are means, with standard deviations in parentheses. IFG = inferior frontal gyrus; SD = standard deviation.

###### Association Condition

Error rates in the association condition were analysed using a 2 × 2 × 2 mixed-design ANOVA with the factors Stimulation (anodal vs sham), Interference (associatively related vs unrelated), and Region (left IFG vs right IFG). The main effect of Stimulation was not significant, *F*(1, 50) = 0.01, *p* = .918, η²1J < .001. There was no Stimulation × Region interaction, *F*(1, 50) = 0.04, *p* = .837, η*²*□ < .001, and no Stimulation × Interference interaction, *F*(1, 50) = 0.06, *p* = .803, η*²*□ = .001. The main effect of Interference did not reach significance, *F*(1, 50) = 2.14, *p* = .150, η*²*□ = .041. However, a significant three-way Stimulation × Interference × Region interaction emerged, *F*(1, 50) = 4.57, *p* = .038, η*²*□ = .084.

To explore this interaction, follow-up 2 × 2 (Stimulation × Interference) ANOVAs were conducted separately for each region. The Stimulation × Interference interaction was not significant in either region (both *p*s > .11). Descriptively, the two regions showed opposite patterns across stimulation conditions (see Figure 3). In the left IFG group, error rates in associatively related trials were lower under anodal than sham stimulation, while error rates in unrelated trials were higher under anodal than sham. In the right IFG group, the pattern was reversed: error rates in associatively related trials were higher under anodal than sham stimulation, while error rates in unrelated trials were lower under anodal than sham.

**Fig. 3.**
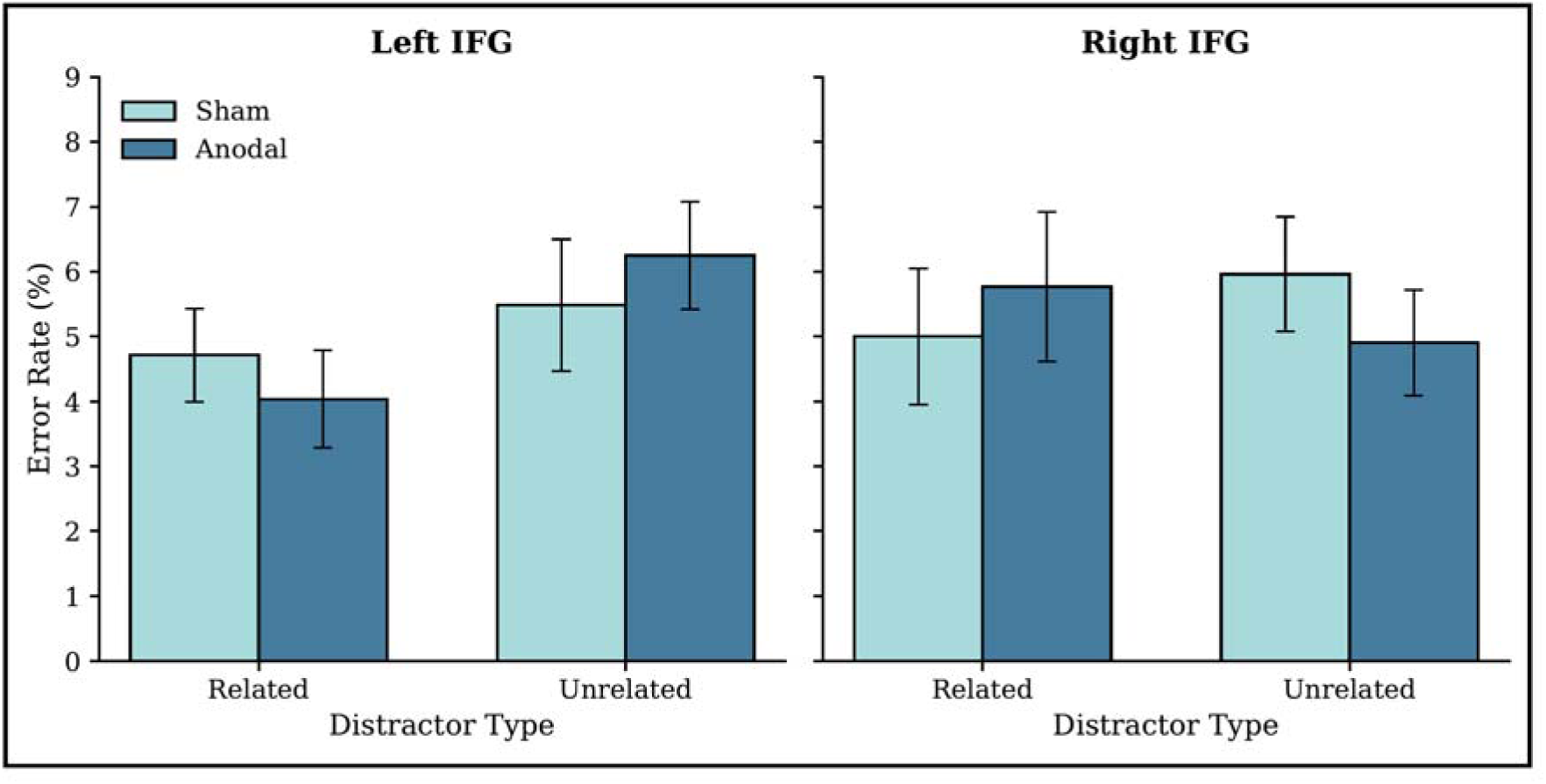
Associative error rates during picture–word interference under sham and anodal stimulation, shown separately for the left IFG and right IFG groups. *Note*. Bars show group means for related and unrelated distractors. Error bars represent the standard error of the mean.

### 3.3. Study 2

#### 3.3.1. Picture Description Task

A target image was presented on the screen. Participants were instructed to describe its contents in full sentences. No time limit was imposed, and participants continued speaking until they decided that they had nothing further to add. Spoken responses were audio-recorded for later transcription. The picture presented in each session was counterbalanced across stimulation conditions, such that half of the participants in each group described one picture under sham and the other under anodal stimulation, while the remaining half received the opposite pairing.

Audio recordings were transcribed using the Descript transcription software (descript.com), and transcripts were manually checked to ensure accuracy. Five connected-speech measures were derived from each transcript: words per minute (WPM), unpruned words per minute (UWPM), syllables per minute (SPM), moving-average type-token ratio (MATTR), and mean length of utterance (MLU). WPM was defined as the count of all intelligible words divided by total speaking time in minutes. Fillers and self-corrections were excluded from this count. UWPM was computed using the same procedure but with fillers and self-corrections retained. SPM was computed as the total number of syllables produced divided by the total speaking time. MATTR was computed over a 50-word moving window following Covington and McFall (2010), and MLU was computed as the mean number of words per T-unit, with a T-unit defined as a main clause together with any subordinate or embedded clauses attached to it (Hunt, 1965). Speech samples with fewer than 50 pruned words were excluded from the analyses, as the 50-word moving window required for MATTR computation could not be applied below this threshold.

#### 3.3.2. Stimuli

Two picture description stimuli were used to elicit connected speech samples: the picnic scene from the Western Aphasia Battery–Revised (WAB-R; Kertesz, 2007) and the updated “Cookie Theft” picture (Berube et al., 2019). Both stimuli have been used extensively in connected speech research with healthy adults and clinical populations, including studies that report speech rate and related microstructural measures (Wilson et al., 2010; Forbes-McKay & Venneri, 2005). The original Cookie Theft picture, included in the Boston Diagnostic Aphasia Examination (Goodglass & Kaplan, 1983), was developed over five decades ago and has since been criticised as anachronistic, depicting gender-stereotyped domestic roles, dated visual content, and culturally narrow imagery that no longer reflects contemporary populations (Steinberg et al., 2022). Berube et al. (2019) introduced a modernised, colour-rendered version of the stimulus to address these limitations. Therefore, the modernised version was adopted in this study.

The WAB-R picnic scene is a black-and-white outdoor depiction with multiple human figures, animals, and objects, whereas the modernised Cookie Theft is a colour-rendered indoor domestic scene. The picture used for elicitation is known to shape the speech sample it produces (Stark, 2019; Schnur & Wang, 2024). Pairing a monochrome outdoor picture with a colour indoor picture was therefore intended to offset two properties that could favour one picture over the other for verbal output. Colour images tend to elicit greater visual engagement than monochrome images, which would advantage the Cookie Theft. Outdoor scenes typically have a broader spatial spread of describable content than indoor ones, which would conversely advantage the WAB picnic.

This pairing rationale was empirically supported by data from a pilot study conducted prior to the main experiment. Speech rate measures were compared between the two stimuli using Mann-Whitney U tests, given the small pilot sample sizes (see Table 5). No significant differences were observed in words per minute or syllables per minute. Effect sizes were small, supporting the comparability of the two stimuli on speech rate measures.

**Table 5.** Comparison of the Picnic Scene and the Updated Cookie Theft Picture on Five Speech Measures.

| Measure | Picnic Scene<br>*M (SD)* | Cookie Theft<br>*M (SD)* | *U* | *p* | *r* |
| --- | --- | --- | --- | --- | --- |
| WPM | 117.31 (18.01) | 113.08 (27.43) | 121 | .499 | .15 |
| UWPM | 126.03 (20.25) | 119.76 (29.13) | 124 | .419 | .18 |
| SPM | 157.17 (22.97) | 151.22 (33.07) | 121 | .499 | .15 |
| MATTR | 0.68 (0.06) | 0.71 (0.05) | 70 | .132 | -.33 |
| MLU | 10.79 (1.99) | 11.23 (2.95) | 97 | .743 | -.08 |

*Note.* Picnic Scene group $n = 14$ ; updated Cookie Theft group $n = 15$ . WPM = words per minute. UWPM = unpruned words per minute. SPM = syllables per minute. MATTR = moving-average type-token ratio. MLU = mean length of utterance. *M* = mean. *SD* = standard deviation. *U* = Mann-Whitney *U* statistic. *r* = rank-biserial correlation effect size.

#### 3.3.3. Results

##### 3.3.3.1. Words per minute

The complete ANOVA output for all speech rate and lexical measures (WPM, UWPM, SPM, MATTR, MLU) is reported in Supplementary Table S2. For WPM, the ANOVA showed a significant Stimulation × Region interaction, *F*(1, 50) = 4.17, *p* = .046, η²p = .077. The main effects of Stimulation, *F*(1, 50) = 3.19, *p* = .080, η*²*□ = .060, and Region, *F*(1, 50) = 0.49, *p* = .487, η*²*□ = .010, were not significant. Follow-up analyses showed that the interaction was driven by the right IFG group, in which WPM was higher under anodal than sham stimulation, *F*(1, 25) = 7.40, *p* = .012, η*²*□ = .228. In the left IFG group, WPM did not differ between the two conditions, *F*(1, 25) = 0.03, *p* = .858, η*²*□ = .001.

##### 3.3.3.2. Unpruned words per minute

The same pattern was found for UWPM. The Stimulation × Region interaction was significant, *F*(1, 50) = 5.45, *p* = .024, η*²*□ = .098, while the main effects of Stimulation, *F*(1, 50) = 3.33, *p* = .074, η*²*□ = .062, and Region, *F*(1, 50) = 0.04, *p* = .845, η*²*□ < .001, were not. Follow-up tests showed that UWPM was higher under anodal than sham stimulation in the right IFG group, *F*(1, 25) = 7.82, *p* = .010, η*²*□ = .238, but did not differ between the conditions in the left IFG group, *F*(1, 25) = 0.15, *p* = .706, η*²*□ = .006.

##### 3.3.3.3. Syllables per minute

The ANOVA for SPM showed a significant main effect of Stimulation, *F*(1, 50) = 4.50, *p* = .039, η*²*□ = .083, and a significant Stimulation × Region interaction, *F*(1, 50) = 4.61, *p* = .037, η*²*□ = .084. The main effect of Region was not significant, *F*(1, 50) < 0.01, *p* = .976, η*²*□ < 0.001. As in the previous two measurements, the interaction was driven by the increase in the right IFG group due to anodal stimulation, *F*(1, 25) = 8.68, *p* = .007, η*²*□ = .258. The left IFG group showed no difference across conditions, *F*(1, 25) < 0.01, *p* = 0.986, η*²*□ < 0.001.

**Fig. 4.**
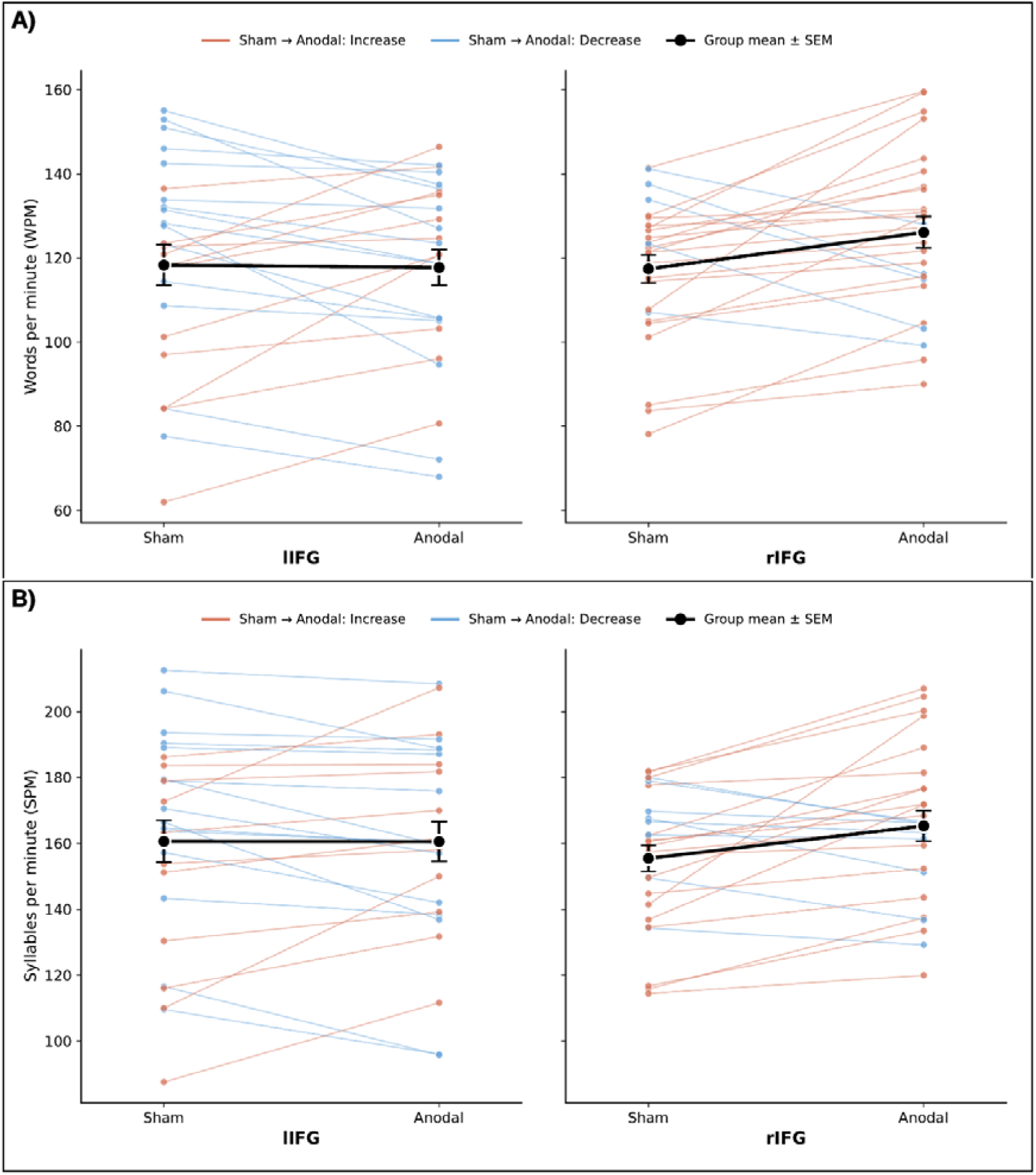
Speech rate during picture description under sham and anodal stimulation, shown separately for the left IFG and right IFG groups. (A) Words per minute (WPM). (B) Syllables per minute (SPM). *Note.* Coloured lines represent individual participants. The black line shows the group mean ± standard error of the mean.

##### 3.3.3.4. Moving-average type-token ratio

MATTR was not affected by either factor. None of the ANOVA effects were significant: Stimulation, *F*(1, 50) = 0.47, *p* = .498, η*²*□ = .009; Region, *F*(1, 50) = 0.03, *p* = .865, η*²*□ < .001; Stimulation × Region, *F*(1, 50) = 0.17, *p* = .687, η*²*□ = .003. Lexical diversity was therefore similar across stimulation conditions and regions.

##### 3.3.3.5. Mean length of utterance

MLU also showed no significant effects. The ANOVA returned non-significant effects of Stimulation, *F*(1, 50) = 0.71, *p* = .403, η*²*□ = .014, Region, *F*(1, 50) = 0.41, *p* = .523, η*²*□ = .008, and their interaction, *F*(1, 50) = 0.07, *p* = .789, η*²*□ = .001. Utterance length did not change with stimulation or region.

**Table 6.**
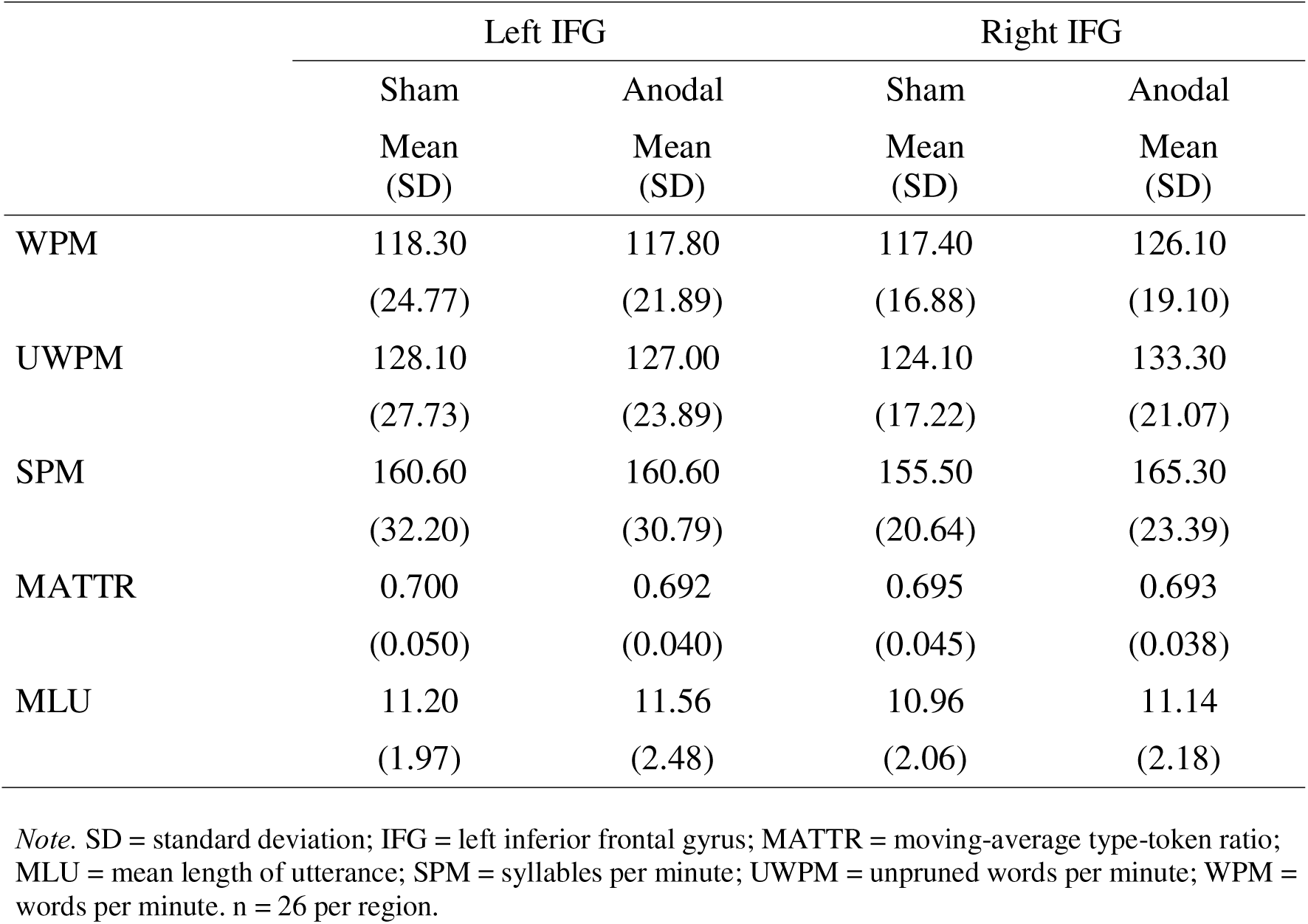
Descriptive statistics of speech rate measures during picture description in the left IFG and right IFG groups.

|  | Left IFG |  | Right IFG |  |
| --- | --- | --- | --- | --- |
|  | Sham<br>Mean<br>(SD) | Anodal<br>Mean<br>(SD) | Sham<br>Mean<br>(SD) | Anodal<br>Mean<br>(SD) |
| WPM | 118.30<br>(24.77) | 117.80<br>(21.89) | 117.40<br>(16.88) | 126.10<br>(19.10) |
| UWPM | 128.10<br>(27.73) | 127.00<br>(23.89) | 124.10<br>(17.22) | 133.30<br>(21.07) |
| SPM | 160.60<br>(32.20) | 160.60<br>(30.79) | 155.50<br>(20.64) | 165.30<br>(23.39) |
| MATTR | 0.700<br>(0.050) | 0.692<br>(0.040) | 0.695<br>(0.045) | 0.693<br>(0.038) |
| MLU | 11.20<br>(1.97) | 11.56<br>(2.48) | 10.96<br>(2.06) | 11.14<br>(2.18) |

*Note.* SD = standard deviation; IFG = left inferior frontal gyrus; MATTR = moving-average type-token ratio; MLU = mean length of utterance; SPM = syllables per minute; UWPM = unpruned words per minute; WPM = words per minute. $n = 26$ per region.

## 4. Discussion

This is the first study to use focal tDCS to examine the dissociable contributions of the left and right inferior frontal gyrus (IFG) to word production in healthy adults. Two distinct paradigms were administered within the same stimulation session. This design provided convergent evidence from constrained naming and connected speech in the same participants. Healthy young adults received anodal and sham stimulation over either the left or right IFG.

Within each session, every participant completed both a picture–word interference task and a picture description task. In the picture–word interference task, semantic interference and associative facilitation emerged reliably in response times but were not modulated by stimulation. However, a significant three-way Stimulation by Interference by Region interaction was observed in error rates for the associative condition, with the two regions showing opposite patterns of modulation. Anodal stimulation of the left IFG enhanced the associative advantage in error rates, whereas anodal stimulation of the right IFG eliminated this benefit and shifted the pattern in the opposite direction. In the picture description task, anodal stimulation of the right IFG increased speech rate, producing significant effects on words per minute, unpruned words per minute, and syllables per minute. No equivalent effects were observed in the left IFG group, and lexical diversity and utterance length were unaffected. Together, these findings provide convergent evidence of a hemispheric dissociation in word production. The right IFG contributes to the regulation of speech output and to the control of distractor interference during constrained naming. The left IFG appears to support lexical-semantic processing.

Our results replicated the standard behavioural signature of the picture–word interference paradigm. Relative to unrelated controls, categorical distractors slowed naming latencies, whereas associative distractors produced faster responses (Schriefers et al., 1990; Alario et al., 2000). However, neither effect was modulated by stimulation in response times. Anodal stimulation of the left IFG did not reduce semantic interference, contrary to our hypothesis. This null pattern is consistent with the mixed picture in the existing tDCS literature on picture naming. Several studies have reported positive effects of stimulation on PWI performance in healthy adults (Wirth et al., 2011; Pisoni et al., 2012), whereas others have failed to find such effects (Westwood et al., 2017). These inconsistencies may stem from differences in stimulation parameters, electrode montage, or stimulus timing across studies.

In the categorical condition, an interesting dissociation emerged between response times and error rates. Naming latencies were slower for categorically related than unrelated distractors, consistent with the classic semantic interference effect. However, error rates showed the opposite pattern, with fewer errors made on related than unrelated trials. This dissociation is best characterised as a speed–accuracy trade-off. Participants appeared to allocate additional processing time to related trials, thereby slowing responses but also reducing naming errors. Such patterns are not uncommon in the PWI literature. They have been interpreted as evidence of strategic response selection in the presence of competing semantic representations (Mahon et al., 2007).

The most striking finding from the PWI task emerged in error rates for the associative condition. The effect of stimulation depended on both the type of interference and the region stimulated. The two regions showed opposite patterns of stimulation effect. Anodal stimulation of the left IFG enhanced the associative advantage, producing fewer errors on related and more errors on unrelated trials. Anodal stimulation of the right IFG produced the reverse pattern, with the associative advantage lost. The follow-up analyses within each region did not reach significance (both ps > .11), but the three-way interaction confirmed that the two regions differed reliably. This opposite pattern is consistent with distinct operations in the two regions. Anodal stimulation of the left IFG may strengthen lexical-semantic processing, which would increase the influence of the distractor word on naming. Anodal stimulation of the right IFG may instead strengthen the suppression of distractor information, reducing its influence (Aron et al., 2014; Nozari & Novick, 2017). On this account, the associative effect in error rates reflects the balance between facilitation and competition during lexical selection (Abdel Rahman & Melinger, 2009). Lemma selection is itself a competitive process in which target and distractor lemmas are activated together (Levelt et al., 1999). Small changes in the balance of activation can change which lemma is selected without necessarily affecting the time taken to complete selection (Roelofs, 1992). Errors may therefore capture lemma-level competition more directly than response times do in this paradigm.

The timing of the associative distractor supports this interpretation. Written distractors reach full lemma activation between 150 and 300 ms after presentation (Hauk et al., 2006; Krott et al., 2019). A distractor presented at SOA −100 ms reaches this peak during the target’s lemma selection window (Indefrey, 2011). Anodal tDCS raises cortical excitability in the targeted region (Stagg & Nitsche, 2011), so its influence on lexical selection should be greatest when distractor activation coincides with this window. The categorical condition showed a different pattern, with semantic interference observed in response times rather than error rates. This contrast may reflect the different SOAs used in the two conditions. The categorical SOA of −300 ms places distractor peak activation near picture onset, producing sustained competition that primarily lengthens response latencies. The associative SOA of −100 ms places distractor peak activation within the lemma selection window itself, where stimulation can shift the outcome of competition without necessarily altering its timing (Levelt et al., 1999; Roelofs, 1992). A different SOA was used in each condition, following the separate literatures for the two effects. The categorical and associative conditions thus differed in SOA and distractor type. The difference in SOA may have contributed to the different stimulation effects observed across the two conditions.

Anodal stimulation of the right IFG increased speech rate during picture description. Significant effects emerged on words per minute, unpruned words per minute, and syllables per minute. The left IFG group showed no equivalent effects. This pattern aligns with models of speech motor control that include a right-lateralised feedback component (Tourville & Guenther, 2011). The right IFG has also been shown to support faster responses during speech production, although that evidence comes from a setting in which the left IFG was disrupted (Hartwigsen et al., 2013). The present finding extends this by showing that directly increasing right IFG excitability can raise speech rate while the left IFG remains intact. Previous tDCS work on connected speech has been conducted largely in clinical populations. These include chronic aphasia (Marangolo et al., 2014), developmental stuttering (Chesters et al., 2018), and progressive supranuclear palsy (Madden et al., 2019). One study in healthy older adults did not find a clear dissociation between the two regions (Matar et al., 2020). The present results provide the first evidence that anodal stimulation of the right IFG can selectively increase speech rate in unimpaired young speakers. By contrast, anodal stimulation of the left IFG did not affect lexical diversity or utterance length, contrary to our prediction. The reasons for this null pattern are unclear, but it may reflect ceiling-level performance on these measures in unimpaired individuals.

Considered together, the two studies provide convergent evidence for a hemispheric dissociation in word production. The right IFG modulated word production in both paradigms. Anodal stimulation of this region increased speech rate in picture description and altered the associative effect on error rates in picture–word interference. Both effects fit a role for the right IFG in the control and regulation of speech output. This view is consistent with accounts that situate the right IFG within a wider system for monitoring and inhibition (Aron et al., 2014; Tourville & Guenther, 2011). The left IFG showed a more limited contribution. Its modulation was confined to error rates in the associative condition, and connected speech measures were not affected by stimulation of this region. This asymmetry is consistent with the view that the right IFG operates as a weakly active node of a bilateral language network. Its contribution becomes detectable when stimulation increases a function that is normally masked by left-hemisphere dominance (Turkeltaub et al., 2011; Hartwigsen & Saur, 2019). The present findings extend this account by showing that the right IFG can also exert a modulatory role in unimpaired speakers when its activity is selectively increased.

Several methodological features support the interpretation of these findings. The focal ring montage used in the present study limits current spread to adjacent cortical regions compared with conventional tDCS (Bortoletto et al., 2016). Stimulation was therefore more closely confined to the targeted area, allowing inferences to be drawn about the left and right IFG with greater anatomical specificity. The double-blind, sham-controlled design minimised expectancy effects. Mood did not differ between stimulation conditions, indicating that the observed effects were not driven by changes in affective state. Adverse effects were generally mild, although they were stronger under anodal stimulation of the right IFG than under sham. This difference did not undermine blinding. Finally, the administration of both paradigms within the same stimulation session allowed convergent inferences to be drawn from the same participants, rather than from separate samples.

The present study has several limitations. First, the order of the two tasks was not counterbalanced. Each participant completed the picture–word interference task before the picture description task in both sessions. This controls for any systematic effect of order at the group level. However, the lack of counterbalancing means that fatigue, practice, or carry-over effects from the preceding PWI task cannot be entirely ruled out. Second, the sample of 52 participants per study was relatively small, which may have limited statistical power. Third, the picture–word interference task used a single SOA for each distractor type. While these SOAs were selected based on separate literatures for categorical and associative effects, the absence of multiple SOAs prevents a direct test of how stimulation effects vary across the time course of lexical access. Future studies systematically varying SOA would help clarify the temporal window in which stimulation-induced modulation of lexical selection is most observable. Fourth, the sample consisted of healthy young adults, which limits generalisation to clinical populations and older speakers. Finally, electrode placement was guided by the international 10–20 EEG system and did not incorporate individual structural imaging.

## 5. Conclusions

In sum, the present study used focal tDCS to test the contributions of the left and right inferior frontal gyrus to word production across two distinct paradigms within the same stimulation session. Anodal stimulation of the right IFG increased speech rate during picture description and modulated the associative effect on error rates during picture–word interference. Anodal stimulation of the left IFG modulated associative error rates in the opposite direction, but did not affect response times or measures of connected speech. Together, these findings provide convergent evidence of a hemispheric dissociation in word production. The right IFG appears to support the regulation of speech output and the control of distractor interference, whereas the left IFG appears to support the processing of word meanings during selection. Future research should extend this dissociation to clinical populations and adopt designs that test its underlying neural mechanisms. Replication in larger samples will also be important to confirm the patterns observed here.

## Supporting information

Statistical Analysis Results

## CRediT authorship contribution statement

**Abdulbaki Yucel:** Conceptualization, Methodology, Formal analysis, Investigation, Visualization, Project administration, Writing – original draft, Writing – review & editing. **Andrew K. Martin:** Conceptualization, Formal analysis, Data curation, Supervision, Project administration, Writing – review & editing.

## Declaration of competing interest

The authors report no conflicts of interest.

## Acknowledgements

The authors are grateful to all participants who took part in this study, and to the Ministry of National Education of the Republic of Türkiye for its support.

## Funding

This work was supported by the YLSY scholarship programme of the Ministry of National Education of the Republic of Türkiye, awarded to the first author.

## Notes

### Competing Interest Statement

The authors have declared no competing interest.

