## Supplementary material for "Hemispheric Dissociation in the Cognitive Control of Word Production": Statistical Analysis Results

**Supplementary Materials**

Hemispheric Dissociation in the Cognitive Control of Word Production: A Focal tDCS Study of the Left and Right Inferior Frontal Gyrus

Supplementary Tables S1–S3 present the full statistical output for the analyses reported in the main text. Tables were generated in JASP. Region: 1 = left IFG, 2 = right IFG. The symbol ✻ denotes an interaction between factors, for example, Stim ✻ Region. Abbreviations are as follows. IFG = inferior frontal gyrus; lIFG = left IFG; rIFG = right IFG; Stim = stimulation; AE = adverse effects; VAMS = Visual Analogue Mood Scale; WPM = words per minute; UWPM = unpruned words per minute; SPM = syllables per minute; MATTR = moving-average type-token ratio; MLU = mean length of utterance; N = number of participants; df = degrees of freedom; SD = standard deviation; SE = standard error; η²p = partial eta squared.

**Supplementary Table S1. Picture–word interference: naming response times and error rates, with the associative error-rate analysis split by region.**

**Categorical**

| *Within Subjects Effects* | | | | | | |
| --- | --- | --- | --- | --- | --- | --- |
| Cases | Sum of Squares | df | Mean Square | F | p | η²_p_ |
| Stim | 0.052 | 1 | 0.052 | 2.738 | .104 | 0.052 |
| Stim ✻ Region | 0.006 | 1 | 0.006 | 0.320 | .574 | 0.006 |
| Residuals | 0.941 | 50 | 0.019 |  |  |  |
| Interference | 0.027 | 1 | 0.027 | 4.962 | .030 | 0.090 |
| Interference ✻ Region | 0.007 | 1 | 0.007 | 1.229 | .273 | 0.024 |
| Residuals | 0.268 | 50 | 0.005 |  |  |  |
| Stim ✻ Interference | 5.538×10^-4^ | 1 | 5.538×10^-4^ | 0.236 | .629 | 0.005 |
| Stim ✻ Interference ✻ Region | 7.120×10^-4^ | 1 | 7.120×10^-4^ | 0.303 | .584 | 0.006 |
| Residuals | 0.117 | 50 | 0.002 |  |  |  |
| *Note.*  Type III Sum of Squares | | | | | | |

| *Between Subjects Effects* | | | | | | |
| --- | --- | --- | --- | --- | --- | --- |
| Cases | Sum of Squares | df | Mean Square | F | p | η²_p_ |
| Region | 0.090 | 1 | 0.090 | 0.943 | .336 | 0.019 |
| Residuals | 4.792 | 50 | 0.096 |  |  |  |
| *Note.*  Type III Sum of Squares | | | | | | |

**Descriptives**

| *Descriptives* | | | | | | | |
| --- | --- | --- | --- | --- | --- | --- | --- |
| Stim | Interference | Region | N | Mean | SD | SE | Coefficient of variation |
| Sham | Related | 1 | 26 | 1.078 | 0.184 | 0.036 | 0.171 |
|  |  | 2 | 26 | 1.032 | 0.207 | 0.041 | 0.201 |
|  | Unrelated | 1 | 26 | 1.037 | 0.163 | 0.032 | 0.158 |
|  |  | 2 | 26 | 1.021 | 0.202 | 0.040 | 0.198 |
| Anodal | Related | 1 | 26 | 1.050 | 0.146 | 0.029 | 0.139 |
|  |  | 2 | 26 | 0.990 | 0.177 | 0.035 | 0.179 |
|  | Unrelated | 1 | 26 | 1.023 | 0.150 | 0.029 | 0.147 |
|  |  | 2 | 26 | 0.978 | 0.159 | 0.031 | 0.163 |

**Associative**

| *Within Subjects Effects* | | | | | | |
| --- | --- | --- | --- | --- | --- | --- |
| Cases | Sum of Squares | df | Mean Square | F | p | η²_p_ |
| Stim | 0.027 | 1 | 0.027 | 1.376 | .246 | 0.027 |
| Stim ✻ Region | 5.312×10^-4^ | 1 | 5.312×10^-4^ | 0.027 | .870 | 5.424×10^-4^ |
| Residuals | 0.979 | 50 | 0.020 |  |  |  |
| Interference | 0.059 | 1 | 0.059 | 12.10 | .001 | 0.195 |
| Interference ✻ Region | 0.005 | 1 | 0.005 | 0.935 | .338 | 0.018 |
| Residuals | 0.243 | 50 | 0.005 |  |  |  |
| Stim ✻ Interference | 4.231×10^-4^ | 1 | 4.231×10^-4^ | 0.074 | .787 | 0.001 |
| Stim ✻ Interference ✻ Region | 0.003 | 1 | 0.003 | 0.586 | .447 | 0.012 |
| Residuals | 0.287 | 50 | 0.006 |  |  |  |
| *Note.*  Type III Sum of Squares | | | | | | |

| *Between Subjects Effects* | | | | | | |
| --- | --- | --- | --- | --- | --- | --- |
| Cases | Sum of Squares | df | Mean Square | F | p | η²_p_ |
| Region | 0.085 | 1 | 0.085 | 0.683 | .412 | 0.013 |
| Residuals | 6.207 | 50 | 0.124 |  |  |  |
| *Note.*  Type III Sum of Squares | | | | | | |

**Descriptives**

| *Descriptives* | | | | | | | |
| --- | --- | --- | --- | --- | --- | --- | --- |
| Stim | Interference | Region | N | Mean | SD | SE | Coefficient of variation |
| Sham | Related | 1 | 26 | 1.133 | 0.176 | 0.035 | 0.156 |
|  |  | 2 | 26 | 1.113 | 0.204 | 0.040 | 0.183 |
|  | Unrelated | 1 | 26 | 1.181 | 0.191 | 0.038 | 0.162 |
|  |  | 2 | 26 | 1.126 | 0.210 | 0.041 | 0.187 |
| Anodal | Related | 1 | 26 | 1.118 | 0.148 | 0.029 | 0.132 |
|  |  | 2 | 26 | 1.076 | 0.214 | 0.042 | 0.199 |
|  | Unrelated | 1 | 26 | 1.156 | 0.183 | 0.036 | 0.158 |
|  |  | 2 | 26 | 1.111 | 0.232 | 0.046 | 0.209 |

**Categorical Error Rate**

| *Within Subjects Effects* | | | | | | |
| --- | --- | --- | --- | --- | --- | --- |
| Cases | Sum of Squares | df | Mean Square | F | p | η²_p_ |
| Stim | 0.481 | 1 | 0.481 | 0.024 | .877 | 4.868×10^-4^ |
| Stim ✻ Region | 3.005 | 1 | 3.005 | 0.152 | .698 | 0.003 |
| Residuals | 987.1 | 50 | 19.74 |  |  |  |
| Interference | 254.3 | 1 | 254.3 | 12.86 | < .001 | 0.205 |
| Interference ✻ Region | 34.74 | 1 | 34.74 | 1.756 | .191 | 0.034 |
| Residuals | 989.1 | 50 | 19.78 |  |  |  |
| Stim ✻ Interference | 4.327 | 1 | 4.327 | 0.527 | .471 | 0.010 |
| Stim ✻ Interference ✻ Region | 1.082 | 1 | 1.082 | 0.132 | .718 | 0.003 |
| Residuals | 410.2 | 50 | 8.204 |  |  |  |
| *Note.*  Type III Sum of Squares | | | | | | |

| *Between Subjects Effects* | | | | | | |
| --- | --- | --- | --- | --- | --- | --- |
| Cases | Sum of Squares | df | Mean Square | F | p | η²_p_ |
| Region | 34.74 | 1 | 34.74 | 0.635 | .429 | 0.013 |
| Residuals | 2737 | 50 | 54.74 |  |  |  |
| *Note.*  Type III Sum of Squares | | | | | | |

**Descriptives**

| *Descriptives* | | | | | | | |
| --- | --- | --- | --- | --- | --- | --- | --- |
| Stim | Interference | Region | N | Mean | SD | SE | Coefficient of variation |
| Sham | Related | 1 | 26 | 5.769 | 5.512 | 1.081 | 0.955 |
|  |  | 2 | 26 | 5.385 | 4.165 | 0.817 | 0.773 |
|  | Unrelated | 1 | 26 | 7.308 | 3.737 | 0.733 | 0.511 |
|  |  | 2 | 26 | 8.846 | 5.841 | 1.145 | 0.660 |
| Anodal | Related | 1 | 26 | 5.769 | 5.948 | 1.167 | 1.031 |
|  |  | 2 | 26 | 6.154 | 5.578 | 1.094 | 0.906 |
|  | Unrelated | 1 | 26 | 7.019 | 5.149 | 1.010 | 0.734 |
|  |  | 2 | 26 | 8.750 | 4.016 | 0.788 | 0.459 |

**Associative Error Rate**

| *Within Subjects Effects* | | | | | | |
| --- | --- | --- | --- | --- | --- | --- |
| Cases | Sum of Squares | df | Mean Square | F | p | η²_p_ |
| Stim | 0.120 | 1 | 0.120 | 0.011 | .918 | 2.151×10^-4^ |
| Stim ✻ Region | 0.481 | 1 | 0.481 | 0.043 | .837 | 8.597×10^-4^ |
| Residuals | 558.8 | 50 | 11.18 |  |  |  |
| Interference | 30.77 | 1 | 30.77 | 2.136 | .150 | 0.041 |
| Interference ✻ Region | 27.04 | 1 | 27.04 | 1.877 | .177 | 0.036 |
| Residuals | 720.3 | 50 | 14.41 |  |  |  |
| Stim ✻ Interference | 0.481 | 1 | 0.481 | 0.063 | .803 | 0.001 |
| Stim ✻ Interference ✻ Region | 34.74 | 1 | 34.74 | 4.566 | .038 | 0.084 |
| Residuals | 380.4 | 50 | 7.608 |  |  |  |
| *Note.*  Type III Sum of Squares | | | | | | |

| *Between Subjects Effects* | | | | | | |
| --- | --- | --- | --- | --- | --- | --- |
| Cases | Sum of Squares | df | Mean Square | F | p | η²_p_ |
| Region | 4.327 | 1 | 4.327 | 0.080 | .778 | 0.002 |
| Residuals | 2691 | 50 | 53.81 |  |  |  |
| *Note.*  Type III Sum of Squares | | | | | | |

**Descriptives**

| *Descriptives* | | | | | | | |
| --- | --- | --- | --- | --- | --- | --- | --- |
| Stim | Interference | Region | N | Mean | SD | SE | Coefficient of variation |
| Sham | Related | 1 | 26 | 4.712 | 3.628 | 0.712 | 0.770 |
|  |  | 2 | 26 | 5.000 | 5.339 | 1.047 | 1.068 |
|  | Unrelated | 1 | 26 | 5.481 | 5.197 | 1.019 | 0.948 |
|  |  | 2 | 26 | 5.962 | 4.532 | 0.889 | 0.760 |
| Anodal | Related | 1 | 26 | 4.038 | 3.813 | 0.748 | 0.944 |
|  |  | 2 | 26 | 5.769 | 5.906 | 1.158 | 1.024 |
|  | Unrelated | 1 | 26 | 6.250 | 4.257 | 0.835 | 0.681 |
|  |  | 2 | 26 | 4.904 | 4.152 | 0.814 | 0.847 |

**Associative Error Rate Region (left IFG)**

| *Within Subjects Effects* | | | | | | |
| --- | --- | --- | --- | --- | --- | --- |
| Cases | Sum of Squares | df | Mean Square | F | p | η²_p_ |
| Stim | 0.060 | 1 | 0.060 | 0.008 | .928 | 3.344×10^-4^ |
| Residuals | 179.6 | 25 | 7.185 |  |  |  |
| Interference | 57.75 | 1 | 57.75 | 5.565 | .026 | 0.182 |
| Residuals | 259.4 | 25 | 10.38 |  |  |  |
| Stim ✻ Interference | 13.52 | 1 | 13.52 | 1.892 | .181 | 0.070 |
| Residuals | 178.7 | 25 | 7.147 |  |  |  |
| *Note.*  Type III Sum of Squares | | | | | | |

| *Between Subjects Effects* | | | | | |
| --- | --- | --- | --- | --- | --- |
| Cases | Sum of Squares | df | Mean Square | F | p |
| Residuals | 1203 | 25 | 48.13 |  |  |
| *Note.*  Type III Sum of Squares | | | | | |

**Descriptives**

| *Descriptives* | | | | | | |
| --- | --- | --- | --- | --- | --- | --- |
| Stim | Interference | N | Mean | SD | SE | Coefficient of variation |
| Sham | Related | 26 | 4.712 | 3.628 | 0.712 | 0.770 |
|  | Unrelated | 26 | 5.481 | 5.197 | 1.019 | 0.948 |
| Anodal | Related | 26 | 4.038 | 3.813 | 0.748 | 0.944 |
|  | Unrelated | 26 | 6.250 | 4.257 | 0.835 | 0.681 |

**Associative Error Rate Region (right IFG)**

| *Within Subjects Effects* | | | | | | |
| --- | --- | --- | --- | --- | --- | --- |
| Cases | Sum of Squares | df | Mean Square | F | p | η²_p_ |
| Stim | 0.541 | 1 | 0.541 | 0.036 | .852 | 0.001 |
| Residuals | 379.1 | 25 | 15.17 |  |  |  |
| Interference | 0.060 | 1 | 0.060 | 0.003 | .955 | 1.304×10^-4^ |
| Residuals | 460.9 | 25 | 18.44 |  |  |  |
| Stim ✻ Interference | 21.69 | 1 | 21.69 | 2.688 | .114 | 0.097 |
| Residuals | 201.7 | 25 | 8.070 |  |  |  |
| *Note.*  Type III Sum of Squares | | | | | | |

| *Between Subjects Effects* | | | | | |
| --- | --- | --- | --- | --- | --- |
| Cases | Sum of Squares | df | Mean Square | F | p |
| Residuals | 1487 | 25 | 59.49 |  |  |
| *Note.*  Type III Sum of Squares | | | | | |

**Descriptives**

| *Descriptives* | | | | | | |
| --- | --- | --- | --- | --- | --- | --- |
| Stim | Interference | N | Mean | SD | SE | Coefficient of variation |
| Sham | Related | 26 | 5.000 | 5.339 | 1.047 | 1.068 |
|  | Unrelated | 26 | 5.962 | 4.532 | 0.889 | 0.760 |
| Anodal | Related | 26 | 5.769 | 5.906 | 1.158 | 1.024 |
|  | Unrelated | 26 | 4.904 | 4.152 | 0.814 | 0.847 |

**Supplementary Table S2. Picture description: speech rate and lexical measures (WPM, UWPM, SPM, MATTR, MLU), with region splits.**

**WPM**

| *Within Subjects Effects* | | | | | | |
| --- | --- | --- | --- | --- | --- | --- |
| Cases | Sum of Squares | df | Mean Square | F | p | η²_p_ |
| Stim | 427.3 | 1 | 427.3 | 3.190 | .080 | 0.060 |
| Stim ✻ Region | 558.7 | 1 | 558.7 | 4.171 | .046 | 0.077 |
| Residuals | 6698 | 50 | 134.0 |  |  |  |
| *Note.*  Type III Sum of Squares | | | | | | |

| *Between Subjects Effects* | | | | | | |
| --- | --- | --- | --- | --- | --- | --- |
| Cases | Sum of Squares | df | Mean Square | F | p | η²_p_ |
| Region | 361.1 | 1 | 361.1 | 0.490 | .487 | 0.010 |
| Residuals | 36860 | 50 | 737.3 |  |  |  |
| *Note.*  Type III Sum of Squares | | | | | | |

**Descriptives**

| *Descriptives* | | | | | | |
| --- | --- | --- | --- | --- | --- | --- |
| Stim | Region | N | Mean | SD | SE | Coefficient of variation |
| Sham | 1 | 26 | 118.3 | 24.77 | 4.857 | 0.209 |
|  | 2 | 26 | 117.4 | 16.88 | 3.310 | 0.144 |
| Anodal | 1 | 26 | 117.8 | 21.89 | 4.293 | 0.186 |
|  | 2 | 26 | 126.1 | 19.10 | 3.746 | 0.151 |

**WPM Region lIFG**

| *Within Subjects Effects* | | | | | | |
| --- | --- | --- | --- | --- | --- | --- |
| Cases | Sum of Squares | df | Mean Square | F | p | η²_p_ |
| Stim | 4.398 | 1 | 4.398 | 0.033 | .858 | 0.001 |
| Residuals | 3382 | 25 | 135.3 |  |  |  |
| *Note.*  Type III Sum of Squares | | | | | | |

| *Between Subjects Effects* | | | | | |
| --- | --- | --- | --- | --- | --- |
| Cases | Sum of Squares | df | Mean Square | F | p |
| Residuals | 23940 | 25 | 957.4 |  |  |
| *Note.*  Type III Sum of Squares | | | | | |

**Descriptives**

| *Descriptives* | | | | | |
| --- | --- | --- | --- | --- | --- |
| Stim | N | Mean | SD | SE | Coefficient of variation |
| Sham | 26 | 118.3 | 24.77 | 4.857 | 0.209 |
| Anodal | 26 | 117.8 | 21.89 | 4.293 | 0.186 |

**WPM Region rIFG**

| *Within Subjects Effects* | | | | | | |
| --- | --- | --- | --- | --- | --- | --- |
| Cases | Sum of Squares | df | Mean Square | F | p | η²_p_ |
| Stim | 981.5 | 1 | 981.5 | 7.401 | .012 | 0.228 |
| Residuals | 3316 | 25 | 132.6 |  |  |  |
| *Note.*  Type III Sum of Squares | | | | | | |

| *Between Subjects Effects* | | | | | |
| --- | --- | --- | --- | --- | --- |
| Cases | Sum of Squares | df | Mean Square | F | p |
| Residuals | 12930 | 25 | 517.1 |  |  |
| *Note.*  Type III Sum of Squares | | | | | |

**Descriptives**

| *Descriptives* | | | | | |
| --- | --- | --- | --- | --- | --- |
| Stim | N | Mean | SD | SE | Coefficient of variation |
| Sham | 26 | 117.4 | 16.88 | 3.310 | 0.144 |
| Anodal | 26 | 126.1 | 19.10 | 3.746 | 0.151 |

**UWPM**

| *Within Subjects Effects* | | | | | | |
| --- | --- | --- | --- | --- | --- | --- |
| Cases | Sum of Squares | df | Mean Square | F | p | η²_p_ |
| Stim | 424.4 | 1 | 424.4 | 3.330 | .074 | 0.062 |
| Stim ✻ Region | 694.5 | 1 | 694.5 | 5.448 | .024 | 0.098 |
| Residuals | 6373 | 50 | 127.5 |  |  |  |
| *Note.*  Type III Sum of Squares | | | | | | |

| *Between Subjects Effects* | | | | | | |
| --- | --- | --- | --- | --- | --- | --- |
| Cases | Sum of Squares | df | Mean Square | F | p | η²_p_ |
| Region | 35.19 | 1 | 35.19 | 0.039 | .845 | 7.704×10^-4^ |
| Residuals | 45640 | 50 | 912.7 |  |  |  |
| *Note.*  Type III Sum of Squares | | | | | | |

**Descriptives**

| *Descriptives* | | | | | | |
| --- | --- | --- | --- | --- | --- | --- |
| Stim | Region | N | Mean | SD | SE | Coefficient of variation |
| Sham | 1 | 26 | 128.1 | 27.73 | 5.438 | 0.216 |
|  | 2 | 26 | 124.1 | 17.22 | 3.377 | 0.139 |
| Anodal | 1 | 26 | 127.0 | 23.89 | 4.686 | 0.188 |
|  | 2 | 26 | 133.3 | 21.07 | 4.133 | 0.158 |

**UWPM Region lIFG**

| *Within Subjects Effects* | | | | | | |
| --- | --- | --- | --- | --- | --- | --- |
| Cases | Sum of Squares | df | Mean Square | F | p | η²_p_ |
| Stim | 16.54 | 1 | 16.54 | 0.145 | .706 | 0.006 |
| Residuals | 2848 | 25 | 113.9 |  |  |  |
| *Note.*  Type III Sum of Squares | | | | | | |

| *Between Subjects Effects* | | | | | |
| --- | --- | --- | --- | --- | --- |
| Cases | Sum of Squares | df | Mean Square | F | p |
| Residuals | 30650 | 25 | 1226 |  |  |
| *Note.*  Type III Sum of Squares | | | | | |

**Descriptives**

| *Descriptives* | | | | | |
| --- | --- | --- | --- | --- | --- |
| Stim | N | Mean | SD | SE | Coefficient of variation |
| Sham | 26 | 128.1 | 27.73 | 5.438 | 0.216 |
| Anodal | 26 | 127.0 | 23.89 | 4.686 | 0.188 |

**UWPM Region rIFG**

| *Within Subjects Effects* | | | | | | |
| --- | --- | --- | --- | --- | --- | --- |
| Cases | Sum of Squares | df | Mean Square | F | p | η²_p_ |
| Stim | 1102 | 1 | 1102 | 7.818 | .010 | 0.238 |
| Residuals | 3525 | 25 | 141.0 |  |  |  |
| *Note.*  Type III Sum of Squares | | | | | | |

| *Between Subjects Effects* | | | | | |
| --- | --- | --- | --- | --- | --- |
| Cases | Sum of Squares | df | Mean Square | F | p |
| Residuals | 14990 | 25 | 599.5 |  |  |
| *Note.*  Type III Sum of Squares | | | | | |

**Descriptives**

| *Descriptives* | | | | | |
| --- | --- | --- | --- | --- | --- |
| Stim | N | Mean | SD | SE | Coefficient of variation |
| Sham | 26 | 124.1 | 17.22 | 3.377 | 0.139 |
| Anodal | 26 | 133.3 | 21.07 | 4.133 | 0.158 |

**SPM**

| *Within Subjects Effects* | | | | | | |
| --- | --- | --- | --- | --- | --- | --- |
| Cases | Sum of Squares | df | Mean Square | F | p | η²_p_ |
| Stim | 627.4 | 1 | 627.4 | 4.500 | .039 | 0.083 |
| Stim ✻ Region | 642.1 | 1 | 642.1 | 4.606 | .037 | 0.084 |
| Residuals | 6971 | 50 | 139.4 |  |  |  |
| *Note.*  Type III Sum of Squares | | | | | | |

| *Between Subjects Effects* | | | | | | |
| --- | --- | --- | --- | --- | --- | --- |
| Cases | Sum of Squares | df | Mean Square | F | p | η²_p_ |
| Region | 1.191 | 1 | 1.191 | 8.888×10^-4^ | .976 | 1.777×10^-5^ |
| Residuals | 66980 | 50 | 1340 |  |  |  |
| *Note.*  Type III Sum of Squares | | | | | | |

**Descriptives**

| *Descriptives* | | | | | | |
| --- | --- | --- | --- | --- | --- | --- |
| Stim | Region | N | Mean | SD | SE | Coefficient of variation |
| Sham | 1 | 26 | 160.6 | 32.20 | 6.316 | 0.200 |
|  | 2 | 26 | 155.5 | 20.64 | 4.049 | 0.133 |
| Anodal | 1 | 26 | 160.6 | 30.79 | 6.038 | 0.192 |
|  | 2 | 26 | 165.3 | 23.39 | 4.587 | 0.141 |

**SPM Region lIFG**

| *Within Subjects Effects* | | | | | | |
| --- | --- | --- | --- | --- | --- | --- |
| Cases | Sum of Squares | df | Mean Square | F | p | η²_p_ |
| Stim | 0.043 | 1 | 0.043 | 3.209×10^-4^ | .986 | 1.283×10^-5^ |
| Residuals | 3313 | 25 | 132.5 |  |  |  |
| *Note.*  Type III Sum of Squares | | | | | | |

| *Between Subjects Effects* | | | | | |
| --- | --- | --- | --- | --- | --- |
| Cases | Sum of Squares | df | Mean Square | F | p |
| Residuals | 46310 | 25 | 1852 |  |  |
| *Note.*  Type III Sum of Squares | | | | | |

**Descriptives**

| *Descriptives* | | | | | |
| --- | --- | --- | --- | --- | --- |
| Stim | N | Mean | SD | SE | Coefficient of variation |
| Sham | 26 | 160.6 | 32.20 | 6.316 | 0.200 |
| Anodal | 26 | 160.6 | 30.79 | 6.038 | 0.192 |

**SPM Region rIFG**

| *Within Subjects Effects* | | | | | | |
| --- | --- | --- | --- | --- | --- | --- |
| Cases | Sum of Squares | df | Mean Square | F | p | η²_p_ |
| Stim | 1270 | 1 | 1270 | 8.677 | .007 | 0.258 |
| Residuals | 3658 | 25 | 146.3 |  |  |  |
| *Note.*  Type III Sum of Squares | | | | | | |

| *Between Subjects Effects* | | | | | |
| --- | --- | --- | --- | --- | --- |
| Cases | Sum of Squares | df | Mean Square | F | p |
| Residuals | 20670 | 25 | 826.9 |  |  |
| *Note.*  Type III Sum of Squares | | | | | |

**Descriptives**

| *Descriptives* | | | | | |
| --- | --- | --- | --- | --- | --- |
| Stim | N | Mean | SD | SE | Coefficient of variation |
| Sham | 26 | 155.5 | 20.64 | 4.049 | 0.133 |
| Anodal | 26 | 165.3 | 23.39 | 4.587 | 0.141 |

**MTTR**

| *Within Subjects Effects* | | | | | | |
| --- | --- | --- | --- | --- | --- | --- |
| Cases | Sum of Squares | df | Mean Square | F | p | η²_p_ |
| Stim | 5.684×10^-4^ | 1 | 5.684×10^-4^ | 0.467 | .498 | 0.009 |
| Stim ✻ Region | 2.007×10^-4^ | 1 | 2.007×10^-4^ | 0.165 | .687 | 0.003 |
| Residuals | 0.061 | 50 | 0.001 |  |  |  |
| *Note.*  Type III Sum of Squares | | | | | | |

| *Between Subjects Effects* | | | | | | |
| --- | --- | --- | --- | --- | --- | --- |
| Cases | Sum of Squares | df | Mean Square | F | p | η²_p_ |
| Region | 7.495×10^-5^ | 1 | 7.495×10^-5^ | 0.029 | .865 | 5.816×10^-4^ |
| Residuals | 0.129 | 50 | 0.003 |  |  |  |
| *Note.*  Type III Sum of Squares | | | | | | |

**Descriptives**

| *Descriptives* | | | | | | |
| --- | --- | --- | --- | --- | --- | --- |
| Stim | Region | N | Mean | SD | SE | Coefficient of variation |
| Sham | 1 | 26 | 0.700 | 0.050 | 0.010 | 0.072 |
|  | 2 | 26 | 0.695 | 0.045 | 0.009 | 0.065 |
| Anodal | 1 | 26 | 0.692 | 0.040 | 0.008 | 0.057 |
|  | 2 | 26 | 0.693 | 0.038 | 0.007 | 0.055 |

**MLU**

| *Within Subjects Effects* | | | | | | |
| --- | --- | --- | --- | --- | --- | --- |
| Cases | Sum of Squares | df | Mean Square | F | p | η²_p_ |
| Stim | 1.901 | 1 | 1.901 | 0.710 | .403 | 0.014 |
| Stim ✻ Region | 0.193 | 1 | 0.193 | 0.072 | .789 | 0.001 |
| Residuals | 133.9 | 50 | 2.678 |  |  |  |
| *Note.*  Type III Sum of Squares | | | | | | |

| *Between Subjects Effects* | | | | | | |
| --- | --- | --- | --- | --- | --- | --- |
| Cases | Sum of Squares | df | Mean Square | F | p | η²_p_ |
| Region | 2.837 | 1 | 2.837 | 0.414 | .523 | 0.008 |
| Residuals | 342.4 | 50 | 6.848 |  |  |  |
| *Note.*  Type III Sum of Squares | | | | | | |

**Descriptives**

| *Descriptives* | | | | | | |
| --- | --- | --- | --- | --- | --- | --- |
| Stim | Region | N | Mean | SD | SE | Coefficient of variation |
| Sham | 1 | 26 | 11.20 | 1.971 | 0.386 | 0.176 |
|  | 2 | 26 | 10.96 | 2.063 | 0.405 | 0.188 |
| Anodal | 1 | 26 | 11.56 | 2.482 | 0.487 | 0.215 |
|  | 2 | 26 | 11.14 | 2.180 | 0.428 | 0.196 |

**Supplementary Table S3. Adverse effects(AE), mood (VAMS positive and negative), and blinding.**

**AE**

| *Within Subjects Effects* | | | | | | |
| --- | --- | --- | --- | --- | --- | --- |
| Cases | Sum of Squares | df | Mean Square | F | p | η²_p_ |
| Stim | 22.78 | 1 | 22.78 | 6.905 | .011 | 0.119 |
| Stim ✻ Region | 19.16 | 1 | 19.16 | 5.807 | .020 | 0.102 |
| Residuals | 168.3 | 51 | 3.299 |  |  |  |
| *Note.*  Type III Sum of Squares | | | | | | |

| *Between Subjects Effects* | | | | | | |
| --- | --- | --- | --- | --- | --- | --- |
| Cases | Sum of Squares | df | Mean Square | F | p | η²_p_ |
| Region | 67.55 | 1 | 67.55 | 3.979 | .051 | 0.072 |
| Residuals | 865.8 | 51 | 16.98 |  |  |  |
| *Note.*  Type III Sum of Squares | | | | | | |

**Descriptives**

| *Descriptives* | | | | | | |
| --- | --- | --- | --- | --- | --- | --- |
| Stim | Region | N | Mean | SD | SE | Coefficient of variation |
| Sham | 1 | 26 | 2.846 | 2.361 | 0.463 | 0.830 |
|  | 2 | 27 | 3.593 | 3.555 | 0.684 | 0.989 |
| Anodal | 1 | 26 | 2.923 | 2.348 | 0.461 | 0.803 |
|  | 2 | 27 | 5.370 | 4.059 | 0.781 | 0.756 |

**AE_Region left IFG**

| *Within Subjects Effects* | | | | | | |
| --- | --- | --- | --- | --- | --- | --- |
| Cases | Sum of Squares | df | Mean Square | F | p | η²_p_ |
| Stim | 0.077 | 1 | 0.077 | 0.048 | .828 | 0.002 |
| Residuals | 39.92 | 25 | 1.597 |  |  |  |
| *Note.*  Type III Sum of Squares | | | | | | |

| *Between Subjects Effects* | | | | | |
| --- | --- | --- | --- | --- | --- |
| Cases | Sum of Squares | df | Mean Square | F | p |
| Residuals | 237.3 | 25 | 9.492 |  |  |
| *Note.*  Type III Sum of Squares | | | | | |

**Descriptives**

| *Descriptives* | | | | | |
| --- | --- | --- | --- | --- | --- |
| Stim | N | Mean | SD | SE | Coefficient of variation |
| Sham | 26 | 2.846 | 2.361 | 0.463 | 0.830 |
| Anodal | 26 | 2.923 | 2.348 | 0.461 | 0.803 |

**AE_Region right IFG**

| *Within Subjects Effects* | | | | | | |
| --- | --- | --- | --- | --- | --- | --- |
| Cases | Sum of Squares | df | Mean Square | F | p | η²_p_ |
| Stim | 42.67 | 1 | 42.67 | 8.644 | .007 | 0.250 |
| Residuals | 128.3 | 26 | 4.936 |  |  |  |
| *Note.*  Type III Sum of Squares | | | | | | |

| *Between Subjects Effects* | | | | | |
| --- | --- | --- | --- | --- | --- |
| Cases | Sum of Squares | df | Mean Square | F | p |
| Residuals | 628.5 | 26 | 24.17 |  |  |
| *Note.*  Type III Sum of Squares | | | | | |

**Descriptives**

| *Descriptives* | | | | | |
| --- | --- | --- | --- | --- | --- |
| Stim | N | Mean | SD | SE | Coefficient of variation |
| Sham | 27 | 3.593 | 3.555 | 0.684 | 0.989 |
| Anodal | 27 | 5.370 | 4.059 | 0.781 | 0.756 |

**VAMS_positive**

| *Within Subjects Effects* | | | | | | |
| --- | --- | --- | --- | --- | --- | --- |
| Cases | Sum of Squares | df | Mean Square | F | p | η²_p_ |
| Stim | 39.98 | 1 | 39.98 | 0.097 | .757 | 0.002 |
| Stim ✻ Region | 604.0 | 1 | 604.0 | 1.468 | .232 | 0.029 |
| Residuals | 20170 | 49 | 411.6 |  |  |  |
| *Note.*  Type III Sum of Squares | | | | | | |

| *Between Subjects Effects* | | | | | | |
| --- | --- | --- | --- | --- | --- | --- |
| Cases | Sum of Squares | df | Mean Square | F | p | η²_p_ |
| Region | 1188 | 1 | 1188 | 1.253 | .269 | 0.025 |
| Residuals | 46470 | 49 | 948.3 |  |  |  |
| *Note.*  Type III Sum of Squares | | | | | | |

**Descriptives**

| *Descriptives* | | | | | | |
| --- | --- | --- | --- | --- | --- | --- |
| Stim | Region | N | Mean | SD | SE | Coefficient of variation |
| Sham | 1 | 26 | -6.654 | 22.75 | 4.461 | -3.419 |
|  | 2 | 25 | 5.040 | 35.59 | 7.118 | 7.062 |
| Anodal | 1 | 26 | -3.038 | 23.71 | 4.650 | -7.803 |
|  | 2 | 25 | -1.080 | 19.62 | 3.924 | -18.17 |

**VAMS_negative**

| *Within Subjects Effects* | | | | | | |
| --- | --- | --- | --- | --- | --- | --- |
| Cases | Sum of Squares | df | Mean Square | F | p | η²_p_ |
| Stim | 4455 | 1 | 4455 | 2.346 | .132 | 0.046 |
| Stim ✻ Region | 270.8 | 1 | 270.8 | 0.143 | .707 | 0.003 |
| Residuals | 93050 | 49 | 1899 |  |  |  |
| *Note.*  Type III Sum of Squares | | | | | | |

| *Between Subjects Effects* | | | | | | |
| --- | --- | --- | --- | --- | --- | --- |
| Cases | Sum of Squares | df | Mean Square | F | p | η²_p_ |
| Region | 2684 | 1 | 2684 | 0.855 | .360 | 0.017 |
| Residuals | 153800 | 49 | 3138 |  |  |  |
| *Note.*  Type III Sum of Squares | | | | | | |

**Descriptives**

| *Descriptives* | | | | | | |
| --- | --- | --- | --- | --- | --- | --- |
| Stim | Region | N | Mean | SD | SE | Coefficient of variation |
| Sham | 1 | 26 | -14.08 | 38.60 | 7.569 | -2.742 |
|  | 2 | 25 | -21.08 | 50.96 | 10.19 | -2.418 |
| Anodal | 1 | 26 | -24.04 | 45.09 | 8.843 | -1.876 |
|  | 2 | 25 | -37.56 | 63.38 | 12.68 | -1.687 |

**Blinding_Chi**

| *Contingency Tables* | | | |
| --- | --- | --- | --- |
|  | BlindingCorrect | |  |
| Region | 0 | 1 | Total |
| 1 | 9 | 17 | 26 |
| 2 | 12 | 16 | 28 |
| Total | 21 | 33 | 54 |
| *Note.*  Each cell displays the observed counts | | | |

| *Chi-Squared Tests* | | | |
| --- | --- | --- | --- |
|  | Value | df | p |
| Χ² | 0.385 | 1 | .535 |
| N | 54 |  |  |

| *Nominal* | |
| --- | --- |
|  | Value |
| Phi-coefficient | -0.084 |
| Cramer's V | 0.084 |

**Blinding_Binomial Test**

| *Binomial Test* | | | | | |
| --- | --- | --- | --- | --- | --- |
| Variable | Level | Counts | Total | Proportion | p |
| BlindingCorrect | 0 | 21 | 54 | 0.389 | .134 |
|  | 1 | 33 | 54 | 0.611 | .134 |
| *Note.*  Proportions tested against value: 0.5. | | | | | |
